# Early lineage segregation of primary myotubes from secondary myotubes and adult muscle stem cells

**DOI:** 10.1101/2024.09.13.612840

**Authors:** Gauthier Toulouse, William Jarassier, Valérie Morin, Fabien Le Grand, Christophe Marcelle

## Abstract

Myogenesis in amniotes unfolds through two consecutive waves. The primary myotube lineage is characterized by the expression of slow myosin, sometimes in combination with fast myosin, and may serve as a scaffold for the secondary lineage, which expresses exclusively fast myosin. The embryonic origin of these two lineages, their relationship, and their connection to adult muscle stem cells are unknown. Here, we employed innovative strategies, combining novel TCF-LEF/β-catenin signaling reporters with the precise spatiotemporal control of *in vivo* electroporation in avian embryos, to track limb muscle progenitors from early migration to late fetal stages. Strikingly, we uncovered two distinct progenitor populations co-existing from the earliest stages of limb myogenesis, with specific developmental fates: reporter-positive progenitors exclusively form primary myotubes, while reporter-negative progenitors generate secondary myotubes and adult muscle stem cells. Furthermore, we uncovered a novel function of TCF-LEF/β-catenin signaling in regulating the spatial organization of the primary myotube lineage via CXCR4-mediated control of myoblast migration, likely contributing to its proposed organizing function. By redefining the embryonic origins of these myogenic populations, our findings not only resolve a longstanding question in muscle biology but also provide a crucial molecular entry point for understanding the cellular and molecular underpinnings of muscle fiber type diversity and function.

## Introduction

The early steps of skeletal muscle morphogenesis have been extensively studied, leading to the emergence of detailed models of the cellular and molecular mechanisms regulating the first steps of myogenesis^1–8^. As embryos grow, the technical complexity of analyses significantly increases, likely explaining why our knowledge of later events of myogenesis is more fragmented. One of the lesser-known steps of myogenesis is the generation of so-called "primary" and "secondary" myotubes. Microscopists studying muscle formation in vertebrate (birds and rodents) embryos half a century ago have identified two distinct waves of myogenesis. The first, taking place during the embryonic phase of development, generates what they have named primary myotubes. The second wave, observed in fetal life, results in the emergence of secondary myotubes. Distinct morphological characteristics separate primary and secondary myotubes ^9,10^. Primary myotubes are initially large and extend from tendon to tendon of embryonic muscles. Secondary myotubes are initially small and tightly organized around primary myotube ^11–13^. Such an association led to the widely held but unproven assumption that primary myotubes may serve as an organizing scaffold for the morphogenesis of the secondary lineage ^10^. Primary myotubes form before motoneurons reach the muscle masses, and they develop normally until birth despite denervation or inhibition of neural activity ^14,15^. Secondary myotubes, on the contrary, form after motoneurons have reached the muscle masses and most authors agree that the emergence of secondary myotubes is dependent on innervation ^14–19^. Biochemical properties also distinguish primary and secondary myotubes. In rodents and birds, primary myotubes are characterized by their expression of slow myosin (in variable combination with embryonic/fast myosin), whereas secondary myotubes express exclusively embryonic fast myosin ^20–23^. Crucially, this initial, genetically encoded, fiber type pattern is plastic and modulated throughout life by factors such as innervation, exercise, and hormonal influences ^10^.

Studies on the cellular origin of primary and secondary myotubes have identified three successive waves of myoblast populations: embryonic, fetal and adult (or satellite cell) myoblasts ^10^. Embryonic myoblasts are isolated from muscle masses during the first week of development in chicken and up to the second week in mouse; fetal myoblasts thereafter ^24^. The adult/satellite cell population settles under the myofibers’ basal lamina shortly before hatching or birth and serves as a reserve stem cell population for muscle growth and repair after birth and into adulthood. *In vitro*, embryonic and fetal myoblasts exhibit intrinsic differences in fusion ability, proliferation and response to growth factors ^10,25–29^. Furthermore, transcriptomics analyses of these two populations confirmed that they possess distinct genetic programs ^30,31^.

Nonetheless, the lineage connection between these two populations of myoblasts remains a subject of unresolved matter ^10,32^. Limb grafting in chicken embryos suggested that two successive waves of myogenic progenitors, each with distinct characteristics, migrate into the limb bud to form primary and secondary myotubes ^33^. *In vitro* studies have further complicated the issue, yielding conflicting hypotheses that propose either a single evolving population, transitioning from displaying embryonic to fetal characteristics ^29,30,34^ or two distinct populations present in the limb from the onset of its formation ^10,35,36^.

Here, we have analyzed the developmental path that muscle progenitors of chicken embryos follow from the time of their migration away from the wing somites into the limb bud mesenchyme until late fetal life, shortly before hatching. Using the *in vivo* electroporation technique to specifically target the muscle progenitor population and a transcriptional reporter to monitor TCF-LEF/β-catenin dependent signaling activity in real time, we have discovered that TCF-LEF/β-catenin activation distinguishes positive and negative muscle progenitors within the early growing wing bud. Biochemical analyses showed that the reporter’s activity is restricted to early stage (PAX3^+^/PAX7^+^/MYF5^+^/MYOD^-^) of myoblast differentiation and confined to the early phases of embryonic development, while absent thereafter until hatching. We designed a novel, dynamic Tet-on-based tool (named TCF-Trace) to follow the fate of reporter-positive myogenic progenitors. This demonstrated that all reporter-positive myoblasts readily differentiated into primary myotubes, while the reporter-negative myoblasts gave rise to the late-forming, secondary myofibers, and also to satellite cells. Finally, we demonstrated that TCF-LEF/β-catenin signaling plays a crucial role in the spatial distribution of limb primary muscle progenitors, likely through the transcriptional regulation of the migration-regulating chemokine receptor CXCR4. These discoveries not only address a long-standing question in muscle biology but also provides a crucial molecular gateway for understanding how these early developmental processes contribute to the fiber type diversity of muscle fibers.

## Results

### TCF-LEF/β-catenin dependent signaling is restricted to early limb muscle development

To develop efficient reporters for monitoring the dynamics of signaling pathways, we recently generated avian transgenic lines that revealed unexpected features of TCF/β-catenin-responding cells and tissues. Among those, we were intrigued by the presence of cells dynamically activating the pathway within muscle-forming domains ^37^. To verify the possibility that TCF/β-catenin-dependent signaling is active in the myogenic lineage, we have electroporated this reporter (named 16TF-VNP, see Material and Methods section, Figure 1B) in the lateral border (named the VLL) of forelimb somites (somites 17-21) of E2.5 chicken embryos, from which limb muscle progenitor originate (Figure 1A). This and all electroporated constructs used in this study were cloned into expression vectors containing two Tol2 sequences (T2: transposable elements from medaka fish ^38^) surrounding the entire construct to allow their stable integration in the chicken genome in the presence of transposase, provided by a co-electroporated transposase plasmid (CAGGS-Transposase). Since electroporation leads to the mosaic transfection of the targeted cell population, the reporter was co-electroporated with a ubiquitously-expressed fluorescent marker to identify and analyze electroporated cells individually. We followed the activity of the TCF/β-catenin-responding cells in the myogenic lineage throughout embryonic and fetal development.

**Figure 1:**
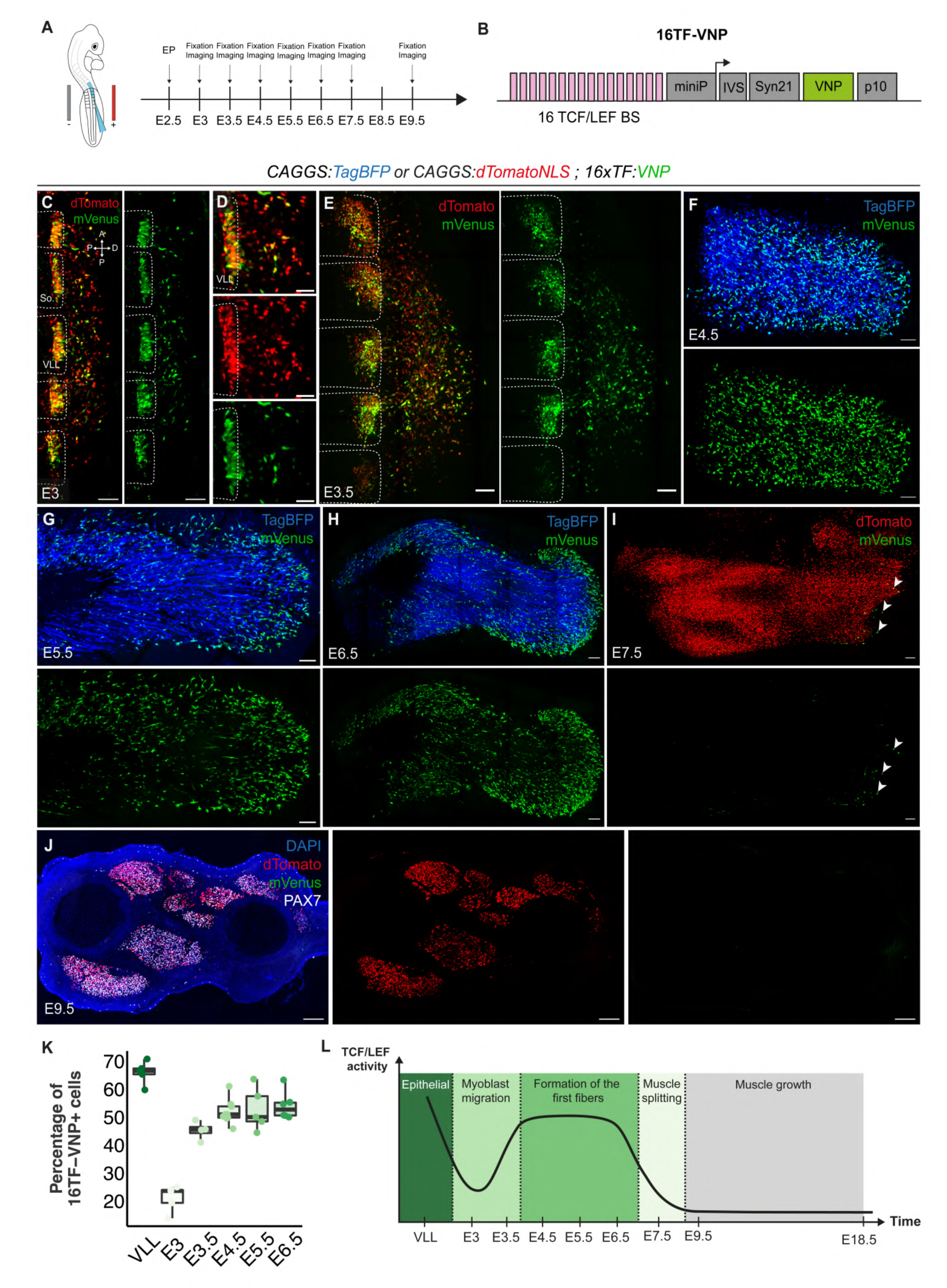
TCF-LEF/β-catenin dependent signaling is restricted to early limb muscle development. **(A)** Brachial somites were electroporated at E2.5 and embryos were analyzed at indicated timepoints **(B)** Representation of the transcriptional reporter (16TF-VNP) used to monitor TCF/LEF/β-catenin dependent signaling. 16 TCF/LEF binding sites (BS) were placed upstream of a minimal promoter (miniP) driving the expression of a nuclear, destabilized Venus fluorescent protein (VNP), three translational enhancers were added (IVS, Syn21 and p10) to boost protein production. **(C-E)** Dorsal view of confocal stacks of brachial somites electroporated with an ubiquitously expressed dTomato and the 16TF-VNP, observed at E3 and E3.5. Somite borders are indicated by dotted lines, **(D)** is an enlargement of **(C)**. **(F-I)** Dorsal views of confocal stacks of limb buds observed between E4.5 and E7.5, electroporated with either an ubiquitous TagBFP **(F,G,H)** or an ubiquitous nuclear dTomato **(I)**, together with the 16TF-VNP. Arrowheads in **(I)** indicate few remaining 16TF-VNP^+^ cells present at E7.5. **(J)** Transversal section of E9.5 limb bud electroporated with an ubiquitous nuclear dTomato together with the 16TF-VNP. **(K)** Quantification of the percentage of 16TF-VNP^+^ cells, between E3 and E6.5. VLL represents the time at which progenitors are located in the epithelial VLL of the brachial somite. **(L)** Schematic representation of 16TF-VNP activity in the myogenic lineage during development. So: somite; VLL: somite ventro-lateral lip; Scale bars: 100μm (C-F) or 200 μm (G-J)

At E3, i.e. twelve hours after electroporation, the migration of progenitors emanating from the VLL has started (Figure 1C,D). At that stage, two distinct locations of TCF/β-catenin-responding cells were observed: a strong expression in a majority (66%) of electroporated epithelial cells located within the VLL and a weaker expression in a minority (21%) of migrating, electroporated cells exiting from the VLL (Figure 1C,K; Supp. Figure 1A,B). The expression of PAX7 in all (16TF-VNP^+^ and 16TF-VNP^-^) electroporated migrating cells confirmed that these are *bona fide* muscle progenitors (Supp. Figure 1A-C). Half a day later, at E3.5, as all muscle progenitors have exited the VLL ^39^, 45% of electroporated muscle progenitors were strongly positive for the 16TF-VNP reporter (Figure 1E,K). During the following three days of development (E4.5 to E6.5) this proportion and level of expression remained relatively stable, in about 50-55% of electroporated cells (Figure 1 F,G,H K). During this time period, 16TF-VNP^+^ cells were distributed among 16TF-VNP^-^cells, with an increasing tendency towards a localization at the distal end of the muscle masses as development proceeded (Figure 1F,G,H). 16TF-VNP^+^ cells were evenly distributed along the dorso-ventral axis of muscles and were also present in the ventral muscle mass of the limb bud (Supp. Figure 1D,E,F). A sharp decrease in the reporter’s activity was however observed on the next day, at E7.5, as 16TF-VNP^+^ cells became sparse and were confined mainly to the distal-most part of the muscle masses (Figure 1I, arrowheads).

We then performed long-term analyses of the reporter’s activity. Embryos electroporated at E2.5 were analyzed on transversal and longitudinal sections of limbs collected at E9.5, E12.5, E14.5, E16.5 and E18.5. Despite a massive labeling of the muscle masses by the ubiquitously expressed electroporation marker, from E9.5 - E18.5, we did not detect any 16TF-VNP^+^ cells at any of the analyzed developmental stages (Figure 1J; Suppl. Figure 2 and Suppl. Figure 3).

These data demonstrate that TCF/LEF transcription in the muscle progenitor population is dynamic (see Figure 1L). An important feature of the reporter’s activity is that it was observed in about 50% of electroporated muscle progenitors from E3.5 until E6.5. During that time period, muscle progenitors differentiated into many multinucleated muscle fibers, the first visible at around E5/E5.5 ^40^. As muscle differentiation progressed and distinct muscle bundles emerged, the reporter activity sharply dropped and was kept silent until hatching.

### TCF-LEF/β-catenin positive cells are PAX3^+^/PAX7^+^/MYF5^+^/MYOD^-^ muscle progenitors

We then investigated the myogenic differentiation state of TCF-LEF/β-catenin activating cells. During myogenesis, muscle progenitors sequentially express different transcription factors that correspond to different phases of myogenic commitment ^8,41,42^. In the mouse and chicken embryos, the proliferative muscle progenitor population comprises PAX7^+^MYF5^-^ slow-dividing and PAX7^+^/MYF5^+^ fast-dividing cells ^43^. MYOD expression signals the exit of progenitors from cell cycle, and MYOG expression corresponds to terminally differentiated, pre-fusing muscle cells ^44^.

In birds, PAX7 and PAX3 proteins are co-expressed in limb myogenic progenitors, from the moment they exit the VLL and migrate into the limb mesenchyme ^45,46^. In fact, PAX3 and PAX7 co-expression persists in all limb myogenic progenitors throughout development, from E10.5 - E16.5, i.e. when progenitors assume satellite cell positions along the myofibers under the basal lamina (Supplementary Figure 4 D-G) ^6^. Therefore, all 16TF-VNP^+^ and 16TF-VNP^-^ progenitors present in the limb at E4.5 co-expressed PAX3 and PAX7 (Supplementary Figure 4 A-C).

To further characterize the molecular and proliferative signature of TCF-LEF/β-catenin activating cells in the limb bud, we performed immunostainings against PAX7, MYF5, MYOD and EdU. At E4.5, all muscle cells present in avian limb muscle masses are mononucleated and express PAX7 ^40^. At this stage, we observed that, while MYF5 expression was widespread throughout the entire muscle progenitor population, MYOD expression was restricted to its central region (Figure 2 A,D). We observed that all 16TF-VNP^+^ cells (100%) expressed MYF5, while only 6% expressed MYOD (Figure 2A-F). At E6.5, many polynucleated MyHC^+^ muscle fibers are present throughout the growing wing and they are tightly intermingled with single-cell progenitors ^40^. Similar to E4.5, we observed at E6.5 that the vast majority of 16TF-VNP^+^ cells (93%) co-expressed PAX7 and MYF5, but that most (85%) 16TF-VNP^+^ did not express MYOD (Figure 2G-J). At E7.5, even though very few progenitors expressed the reporter, all of them were PAX7^+^ (Figure 2K). Even though we have previously shown that all PAX7^+^/MYF5^+^ progenitors are faster-dividing cells than PAX7^+^/MYF5^-^ progenitors ^43^, it was possible that the 16TF-VNP^+^ and 16TF-VNP^-^subpopulations proliferated at different rates. However, labeling of embryos with EdU showed that both the negative and the positive populations displayed the same rate of proliferation (Suppl. Figure 5A-C).

**Figure 2:**
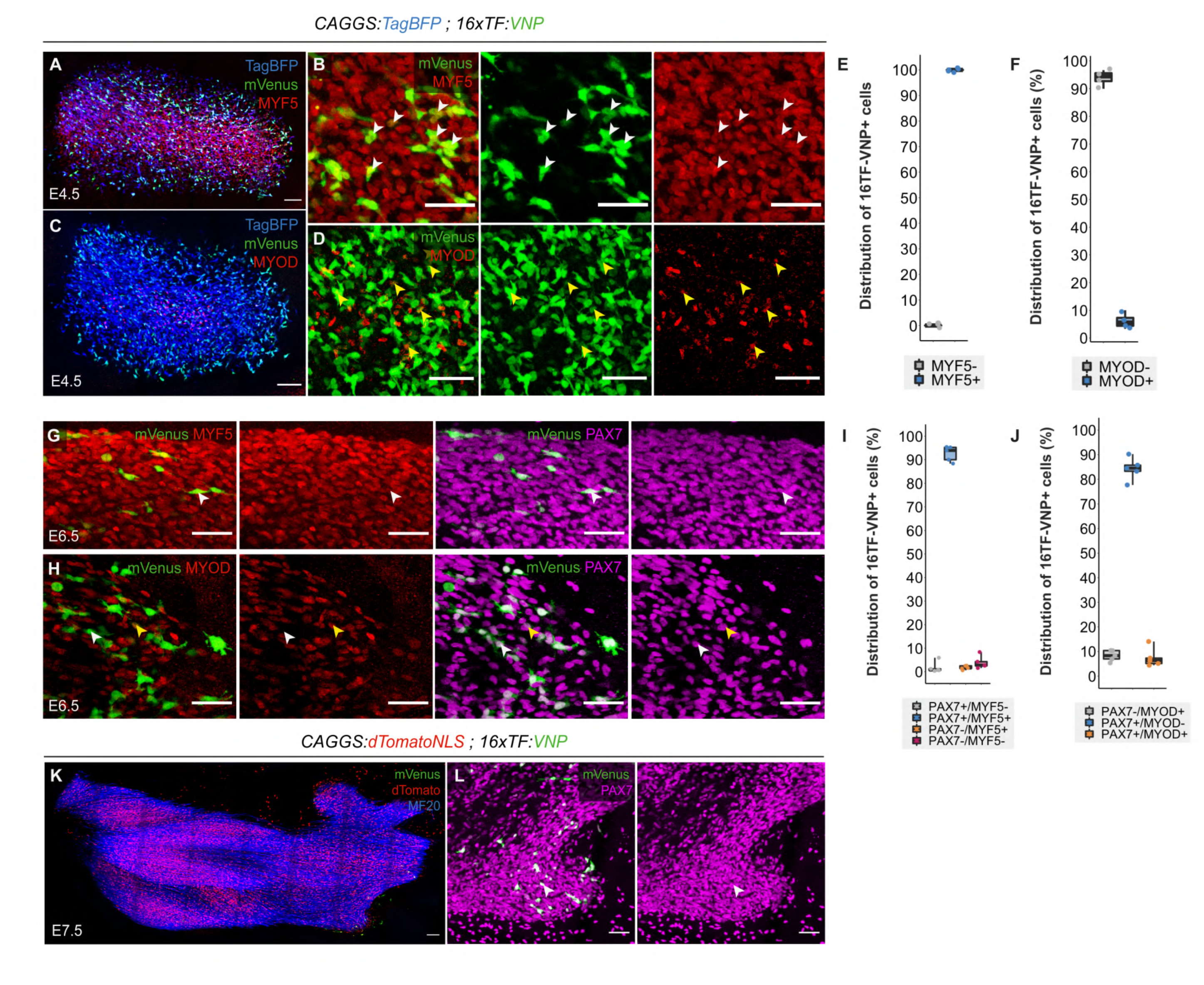
16TF-VNP^+^ cells are early myogenic progenitors. **(A-D)** Dorsal views of confocal stacks of E4.5 limb buds electroporated with an ubiquitous TagBFP (blue), the 16TF-VNP reporter and stained for MYF5 **(A,B)** or MYOD **(C,D).** White arrowheads in **(B)** indicates 16TF-VNP^+^/MYF5^+^ cells; yellow arrowheads in **(D)** indicates some of the few 16TF-VNP^+^/MYOD^+^ observed **(E,F**). Quantification of the percentage of 16TF-VNP^+^ cells positive for MYF5 **(E)** and MYOD **(F). (G,H)** Dorsal view of confocal stacks of E6.5 limb buds electroporated with an ubiquitous TagBFP, the 16TF-VNP reporter and stained for PAX7 and MYF5 **(G)** or PAX7 and MYOD **(H).** The TagBFP channel is not represented. The white arrowheads indicate 16TF-VNP^+^/PAX7^+^/MYF5^+^ **(G)** cell and 16TF-VNP^+^/PAX7^+^/MYOD^-^ **(H)**. The yellow arrowhead in **(H)** indicates a 16TF-VNP^+^/PAX7^+^/MYOD^+^ cell. **(I,J)** Quantification of the percentage of 16TF-VNP^+^ cells positive for PAX7 and MYF5 **(I)** or PAX7 and MYOD **(J). (K,L)** Dorsal view of confocal stacks of E7.5 limb buds electroporated with an ubiquitous nuclear dTomato, the 16TF-VNP reporter and stained for PAX7. The arrowhead indicates one of the few remaining 16TF-VNP^+^ cells that also expresses PAX7. **(L)** is an enlargement of **(K).** Scale bars: 50μm (G-H, L), 100μm (A-D), or 200 μm (K).

Together, these experiments demonstrate that TCF-LEF/β-catenin dependent signaling is strictly restricted to a narrow time window of myogenesis, to a population of proliferating PAX3^+^/PAX7^+^/MYF5^+^ progenitors. As soon as these progenitors progress further along the myogenic path, initiating MYOD expression, and exiting cell cycle, TCF-LEF/β-catenin dependent signaling is turned off.

### TCF-Trace, a dynamic lineage tracing system to follow the fate of TCF-LEF/β-catenin^+^ myogenic precursors

The 16TF-VNP reporter we engineered provides a snapshot of TCF-LEF/β-catenin dependent signaling at the time of analysis. It was however possible that limb muscle progenitors fluctuate between a 16TF-VNP^+^ and a 16TF-VNP^-^ state. To test this, we designed a reporter construct where the destabilized nuclear mVenus fluorescent protein was placed in tandem with a stable nuclear mCherry (half-life: about 18 hours ^47^). This technique has been used in *Drosophilia* ^48^ to test whether the activity of a promoter is fluctuating between an ON/OFF state. If the activity of the reporter was fluctuating, there would be more cells labeled with the stable mCherry than the destabilized mVenus. On the contrary if cells constantly respond to the signal, only double-positive cells should be observed. The construct was electroporated in the VLL at E2.5 and examined in the limb at E4.5. We observed a near-perfect (98%) correlation of mVenus and mCherry stainings, indicating that, within the time frame of the experiment, the 16TF-VNP+ progenitors maintain the reporter’s activity and do not fluctuate between a positive and negative state (Suppl. Figure 6A-C).

It therefore became important to investigate the long-term fate of myogenic progenitors that activate TCF-LEF/β-catenin dependent signaling. To address this, we developed a lineage tracing system using both Tet-On and Cre-Lox technologies (Figure 3 A). This system aims to permanently label cells with dTomato fluorescence when they simultaneously experience TCF-LEF/β-catenin signaling and are exposed to doxycycline (Figure 3B). The Tet-on technology ^49,50^ is a well-established system of drug-induced gene activation, which displays low background and high induction rates, in particular when using the "rtTA/TRE 3G" system ^51,52^. Combining this to the CRE-mediated excision of "Stop/PolyA" sequences placed upstream of a fluorescent protein should deliver a very sensitive system to permanently label myogenic progenitors. However, a significant drawback of the Tet-on system is that the rtTA protein is very stable ^53^, making it unsuitable for dynamic studies. We hypothesized that the rtTA protein could be destabilized, a process that should significantly enhances its utility for dynamic studies but that comes at the cost of significantly weaker protein expression ^48^. We first engineered a rtTA construct where a PEST proteolytic signal was inserted at its N terminus (PEST-rtTA) ^54^.

**Figure 3:**
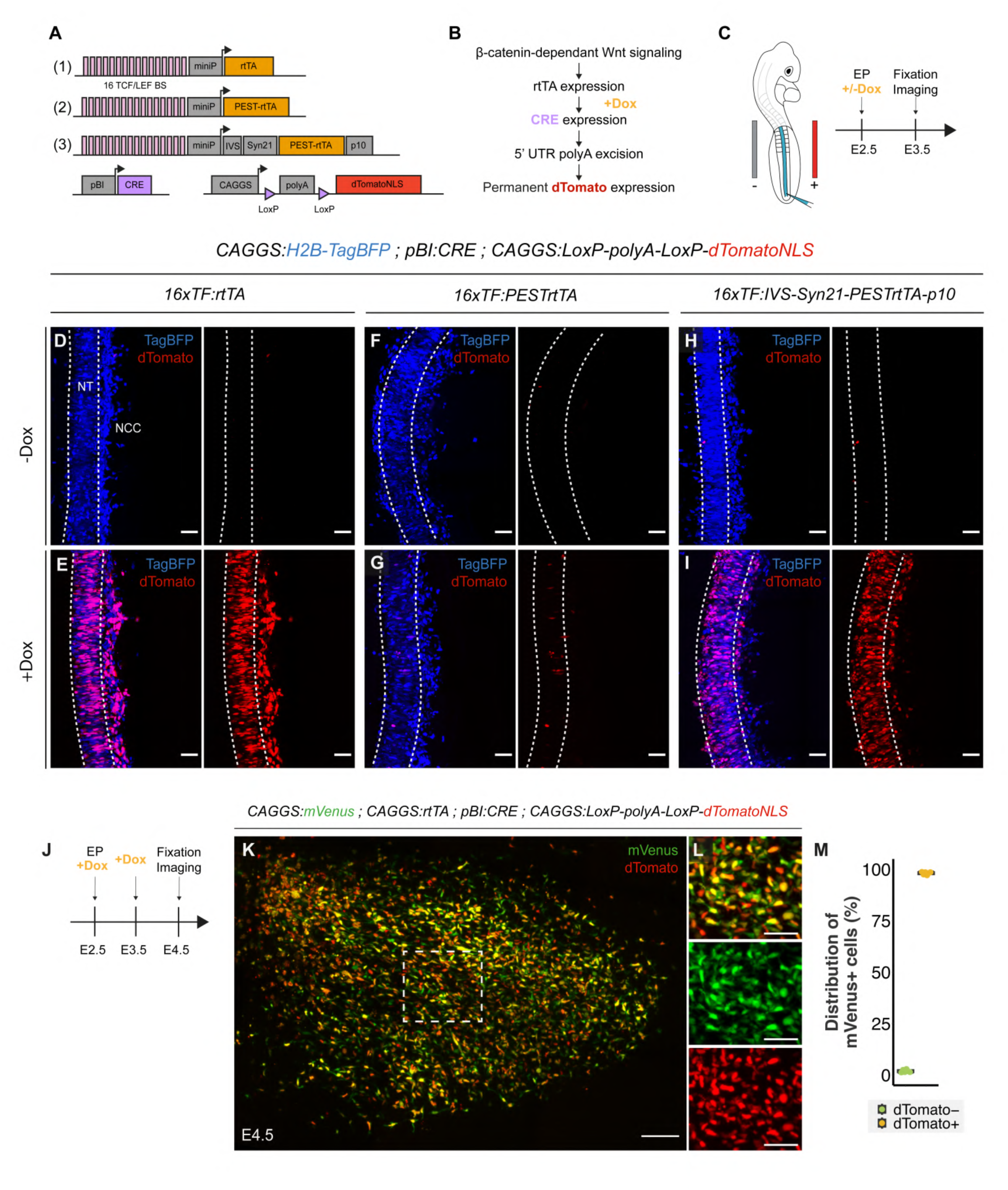
TCF-Trace, a tool to follow the fate of cells experiencing TCF-LEF/β-catenin dependent signaling. **(A,B)** Schematics of the constructs tested. The three constructs comprise 16 TCF/LEF binding sites upstream of a minimal promoter driving the expression of a rtTA (1) or a rtTA fused with a PEST sequence at its N-terminal part (2) or a rtTA fused with a PEST sequence at its N-terminal part, flanked by translational enhancers (3). All constructs were co-electroporated with a plasmid containing a rtTA-dependent CRE recombinase and another containing a CRE-inducible nuclear dTomato. **(C)** Embryos were electroporated in the neural tube at E2.5, induced with doxycycline and analyzed one day later. **(D-I)** Dorsal view of confocal stacks of E3.5 neural tube (NT) electroporated with a ubiquitous H2B-TagBFP and the rtTA plasmid **(D,E)**, the destabilized rtTA **(F,G)** or the destabilized and boosted rtTA **(H,I).** The dotted lines delineate the electroporated, right side of the neural tube. **(J,K)** Dorsal view of confocal stacks of E4.5 limb buds electroporated with an ubiquitous mVenus, an ubiquitous rtTA, the CRE and dTomato plasmids, doxycycline was added at E2.5 and E3.5. **(L)** Quantification of the percentage of dTomato+ cells within the mVenus^+^ population. Each dot represents a limb bud. Scale bars: 50μm (D-I, L) or 100μm (K).

We used the electroporation technique to test this constructs in the dorsal region of the neural tube (Figure 3C), known to respond to TCF/LEF signaling ^37^. In the control experiment, we tested a native (not destabilized) rtTA construct, and we observed that the addition of doxycycline to electroporated embryos led to a strong response, with many visible dTomato^+^ neural cells (Figure 3A,D,E). Expectedly, fusing PEST sequences to the rtTA construct led to a strong decrease in the efficiency of the response, with only few visible dTomato^+^ neural cells (Figure 3A,F,G). To address the reduced expression levels, we incorporated to the rtTA/PEST construct translation enhancer sequences (IVS/Syn21/p10; see above). The addition of translational enhancers to the rtTA-PEST sequence restored a labeling efficiency that was comparable to that observed with the original rtTA construct (Figure 3A,H-I). This suggests that the two-step strategy (destabilization/translation enhancement) generated a sensitive Tet-on system that dynamically responds to doxycycline exposure when TCF-LEF/β-catenin dependent signaling is active.

We tested the efficiency of this tracing system. Tracing TCF-LEF/β-catenin-responding cells involves a succession of molecular steps (doxycycline-triggered activation of Cre expression, excision of Stop sequences and expression of the lineage tracing fluorescent protein, Figure 3B). To evaluate the efficiency of the system to permanently label all cells experiencing TCF-LEF/β-catenin dependent signaling, we substituted the 16TF promoter with a CAGGS ubiquitous promoter. This should theoretically lead to the activation of the tracing fluorescent protein in all electroporated cells upon doxycycline addition. The VLL of brachial somites were co-electroporated with this plasmid mix and embryos were exposed to a doxycycline solution for two consecutive days and then analyzed one day later at E4.5 (Figure 3J). We observed that 98% of the electroporated cells, labeled by the expression of the constitutive mVenus protein, also expressed dTomato (Figure 3K-M). This near-perfect correlation between the expression of mVenus and dTomato suggests that, despite significant destabilization of the rtTA protein, and the many molecular steps required to activate the reporter, the doxycycline-mediated induction of dTomato fluorescence by the Tet-on/CRE system we designed is highly efficient. Furthermore, this experiment indicates that despite using multiple independent plasmids (five, including the transposase construct), all electroporated cells appear to have simultaneously incorporated them. This efficient and dynamic tool, that we named *<u>TCF-Trace</u>*, is the first molecular tool aimed at following the fate of cells experiencing temporary bursts of TCF-LEF/β-catenin-dependent signaling. We therefore proceeded to investigate the fate of myogenic precursor cells labeled by TCF-Trace.

### Two distinct progenitor populations co-exist in early limb myogenesis

The TCF-Trace system displays accurate temporal labeling of targeted cells without significant lag from previous cell history. This temporal precision is crucial in our experimental design, as we electroporate VLL cells that display a high TCF-LEF/β-catenin-dependent activity at E2.5 (Figure 1C-D), an activity, likely linked to their epithelial state ^55–57^, that we do not intend to trace. We therefore initiated the lineage fate of PAX7+/MYF5+ progenitors present in the limb from embryonic day 4.5 (E4.5).

To do this, we electroporated the VLL of E2.5 brachial somites with a combination of the three plasmids described above, together with a ubiquitously expressed mVenus as an electroporation marker and the transposase plasmid. Subsequently, at E4.5, 5.5 and 6.5, doxycycline was added to developing embryos, aiming to label all progenitors activating TCF-LEF/β-catenin-dependent signaling during that time window. We analyzed the embryos at E7.5, at a time when the activity of the 16TF-VNP reporter is almost extinguished (Figure 1L, Figure 4A). We tested for expression of PAX7, mVenus and dTomato proteins (Figure 4B-D). We observed numerous electroporated (green) myofibers containing dTomato^+^ nuclei (red), indicating a massive contribution of the TCF-LEF/β-catenin^+^ progenitors to myotube formation (Figure 4B).

**Figure 4:**
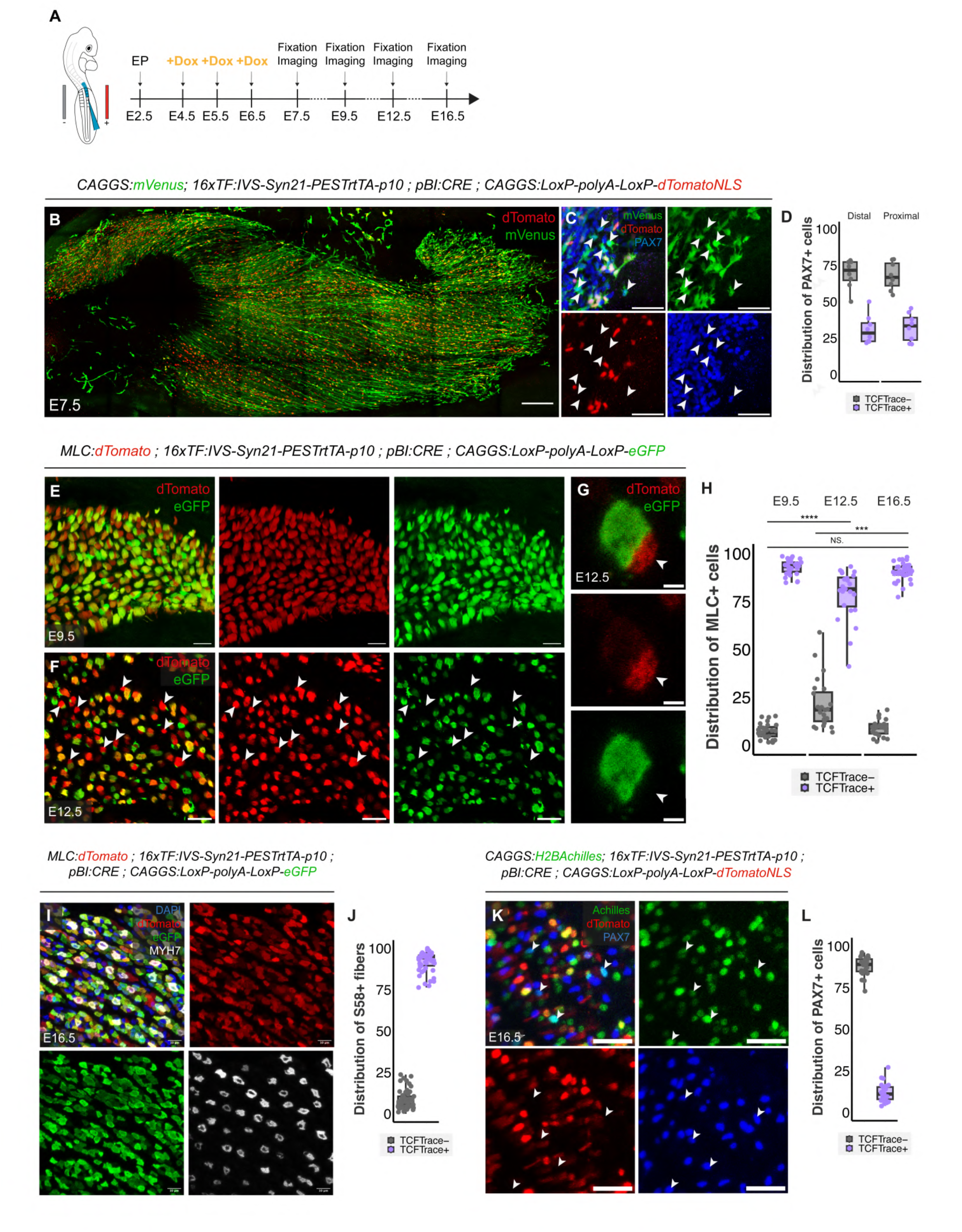
Lineage analysis of TCF-Trace^+^ and TCF-Trace^-^ populations. **(A,B)** TCF-Trace was induced with doxycycline during the time period (E4.5-E6.5) in which muscle progenitors respond to TCF/LEF signaling. Brachial somites of E2.5 embryos were electroporated, TCF-Trace was induced 2-4 days later and embryos were analyzed at various stages from E7.5 to E16.5. **(C-D**) Dorsal view of confocal stacks of E7.5 limb bud electroporated with a ubiquitous mVenus, the TCF-Trace lineage tool driving the expression of a nuclear dTomato and stained for PAX7. **(D)** is an enlargement of **(C). (E)** Quantification of the percentage of TCF-Trace^-^ or TCF-Trace^+^ cells in the PAX7+ electroporated population at E7.5 in the proximal and the distal part of the muscle mass. **(F,G)** Transverse sections of E9.5 **(F**) and E12.5 **(G)** limb buds electroporated with a myofiber-specific dTomato and the TCF-Trace lineage tool driving the expression of eGFP. **(H)** Representative example of a TCF-Trace^+^ myotube surrounded by a smaller TCF-Trace^-^ myotube (white arrowhead). **(I)** Quantification of the percentage of TCF-Trace^-^ and TCF-Trace^+^ myotubes in the MLC^+^ electroporated population. **(J)** Transverse sections of E16.5 limb buds electroporated with a myofiber-specific dTomato, the TCF-Trace lineage tool driving the expression of eGFP and stained for MYH7 (S58 antibody, recognizes slow myosin). **(K)** Quantification of the percentage of TCF-Trace^-^ or TCF-Trace^+^ myotubes in the S58^+^ electroporated myotubes. **(L)** Transverse sections of E16.5 limb buds electroporated with a nuclear Achilles, the TCF-Trace lineage tool driving the expression of a nuclear dTomato and stained for PAX7. **(M)** Quantification of the percentage of TCF-Trace^-^ or TCF-Trace^+^ cells in the PAX7^+^ electroporated population. Each dot represents a section, n=7 limbs. Scale bars: 10μm (J,L,H), 20μm (F,G), 50μm (D), or 200μm (C).

Strikingly, this analysis unveiled an additional finding: a significant proportion (65-72%) of PAX7^+^, electroporated progenitors were not labeled by TCF-Trace. This was observed both in the proximal and distal region of the muscle masses (arrows in Figure 4C, Figure 4D). As we demonstrated the high sensitivity of the tracing system we designed (Figure 3J-M), it is unlikely that the absence of label is due to a failure to detect and trace all TCF-LEF/β-catenin^+^ cells. Instead, it suggests the presence of a population of progenitors in the developing limb that never activate TCF-LEF/β-catenin signaling, co-existing with the TCF-Trace^+^ progenitor population. This intriguing observation triggered further investigation into the long-term fate of these distinct populations.

### TCF-LEF/β-catenin^+^ myogenic precursors differentiate into primary myotubes

To streamline myotube analyses, we engineered a second version of TCF-Trace, where a cytoplasmic forms of the fluorescent protein eGFP was used. This enabled straightforward identification of the entire myotube diameters on cross sections. The electroporation tracer was a cytoplasmic form of dTomato driven by the Myosin Light Chain (MLC ^58^) promoter, known to be expressed in myofibers. On cross sections at E9.5, we observed that nearly all (93%) electroporated myofibers (in red) were positive for eGFP (in green; Figure 4E,H). At this stage in chicken embryos, only primary myotubes are present in muscle masses ^10^. Based on this temporal criterion, we therefore hypothesized that the TCF-LEF/β-catenin^+^ population represents the precursors of primary myotubes.

To confirm this, we proceeded with an immunostaining approach. It is widely accepted that all slow myosin-expressing myofibers are primary myotubes ^20–23^. As reports on slow myosin expression in chicken are scattered over many publications and developmental stages, we repeated a comprehensive survey of slow myosin expression in the chicken forelimb from embryonic to fetal stages of development (Supplementary Figure 7), and observed that its expression significantly progresses along development, starting to be visible at E9.5 in a few of the muscle masses present at that stage (Supplementary Figure 7C), and steadily intensifying to encompass all muscles at E18.5 (Supplementary Figure 7D-H). From E12.5 until E18.5, slow (primary) myofibers are progressively surrounded by small, fast (presumably secondary) myofibers (Supplementary Figure 7I-P). To investigate the slow vs fast signature of the TCF-LEF/β-catenin-derived myofibers during embryonic and fetal development, we performed a lineage tracing experiment similar to the one above but analyzed the embryos at stage E16.5 (Figure 4I-J). We observed that nearly all (98%) slow myosin-positive myotubes expressing the electroporation marker derived from the TCF-LEF/β-catenin^+^-derived lineage (Fig4 I,J). Therefore, based on temporal and biochemical criteria, these experiments demonstrate that TCF-LEF/β-catenin^+^ progenitors present in the growing limb bud constitute the cellular origin of primary myotubes.

### TCF-LEF/β-catenin^-^ myogenic precursors differentiate into secondary myotubes and satellite cells

We then wondered what is the origin of secondary myotubes. Secondary myotubes have been shown to appear after E9 in chicken, often surrounding primary myotubes (Supplementary Figure 7 I,J) ^10,59^. We followed the fate of TCF-Trace-positive and -negative progenitors, using the same protocol as in the previous experiment set, but left the embryos to develop until E12.5. In contrast to the analyses done at E9.5, we found that 21% of electroporated (red) myotubes were not labeled by the TCF-Trace system (green Figure 4F-H). Often, we observed that dTomato-only myotubes were smaller than eGFP-positive myotubes and that they were located at their periphery, which are typical characteristics of secondary myotubes (Figure 4G, arrowheads). Therefore, based on temporal and morphological criteria, these experiments strongly suggest that TCF-LEF/β-catenin^-^ progenitors present in the growing limb bud constitute the cellular origin of secondary myotubes. Interestingly, in a similar analysis done at E16.5, we observed a significant decrease in the proportion of TCF-Trace-negative myotubes, paralleled by an increase in the proportion of TCF-Trace positive myotubes (Figure 4H). It is possible that the shift in the proportions of TCF-Trace^+^ and TCF-Trace^-^ myotubes from E12.5 until E16.5 is due to a mixing between both lineages, either through myoblast to myofibers fusion or through fusion between myofibers.

Finally, we determined whether satellite cells originate from one, the other, or both progenitor populations. VLL cells of E2.5 brachial somites were electroporated with a combination of the three plasmids described above, together with a ubiquitously expressed H2B-Achilles as an electroporation marker. At E4.5, E5.5 and E6.5, doxycycline was added to developing embryos. In birds and rodent, satellite cells are the only PAX7^+^ cells present in muscle masses in late fetal stages, just before hatching/birth ^6,60–63^. We explored whether PAX7^+^ cells present at E16.5 were labelled by the tracing system. We found that the vast majority (87%) of electroporated satellite cells present in limb muscle masses were TCF-Trace-negative, while a minority (13%) was positive (Figure 4K-L). This finding suggests that most satellite cells present in muscles at hatching derive from the TCF-LEF/β-catenin^-^ progenitor population present in early limb buds, with a minor contribution originating from the TCF-LEF/β-catenin^+^ lineage.

### TCF-LEF/β-catenin signaling regulates the spatial distribution of primary progenitors through CXCR4 signaling

With the TCF-LEF/β-catenin reporter revealing pivotal connections between the embryonic origins of primary and secondary myotube lineages and adult muscle stem cells, it naturally prompted us to investigate how this signaling pathway influences these critical developmental processes. Canonical, β-catenin-dependent Wnt signaling has been proposed to fulfill various functions during embryonic myogenesis, synergizing or antagonizing with other signaling pathways, such as Sonic Hedgehog, Bone Morphogenic Proteins, or NOTCH ^42,64–67^. In the limb, conflicting data suggest that β-catenin-dependent Wnt signaling activates, inhibits or is dispensable for embryonic myogenesis ^68–71^. Our discovery of the existence of two co-existing populations of limb muscle progenitors, distinguished by their response to TCF-LEF/β-catenin-dependent signaling, necessitates a reevaluation of the role of Wnt signaling in limb myogenesis. The precise spatio-temporal control provided by *in vivo* electroporation offers a powerful alternative to classical approaches, potentially clarifying the discrepancies observed in previous studies.

To address the role of Wnt β-catenin-dependent signaling in the TCF-LEF/β-catenin-positive primary myotube progenitor population, we used a dominant-negative form of Lef1 (DN Lef1), known to efficiently inhibit Wnt β-catenin-dependent signaling ^72^. This construct was electroporated at E2.5 in the lateral portion of brachial somites, to target the limb muscle progenitor population. Its effect was evaluated at E4.5, at a time when the 16TF-VNP reporter is active in 50% of the progenitor population (Figure 1K-L). We examined myogenic progenitor proliferation and differentiation under this experimental setting. No differences were observed in the progenitor proliferation, as determined by phospho-histone H3 or EdU labeling (Supplementary Figure 8A-F). Similarly, the myogenic differentiation program remained unaltered, since the expression of early differentiation marker MYF5 and of the terminal differentiation marker MYOG were similar to controls (Supplementary Figure 8G-M).

However, the spatial distribution of electroporated cells revealed a notable effect. A subset of these cells exhibited a tendency to remain in a proximal location, closer to the somites from which they originated. In contrast, those that migrated distally showed an increased dispersion away from the limb axis. Quantitative analysis confirmed a significant difference in distribution between the progenitors expressing DN Lef1 and the controls (Figure 5A-D). To rule out the possibility that this phenotype was due to the inhibition of TCF-LEF/β-catenin signaling in the VLL, where it likely maintains its epithelialization state ^55–57^, we repeated the experiment, activating the DN Lef1 expression only after progenitors had migrated out of the VLL and examined embryos at E6.5. The results were consistent with the previous experiment (Supplementary Figure 8N-O), reinforcing the role of TCF-LEF/β-catenin signaling in the control of muscle progenitor distribution.

**Figure 5:**
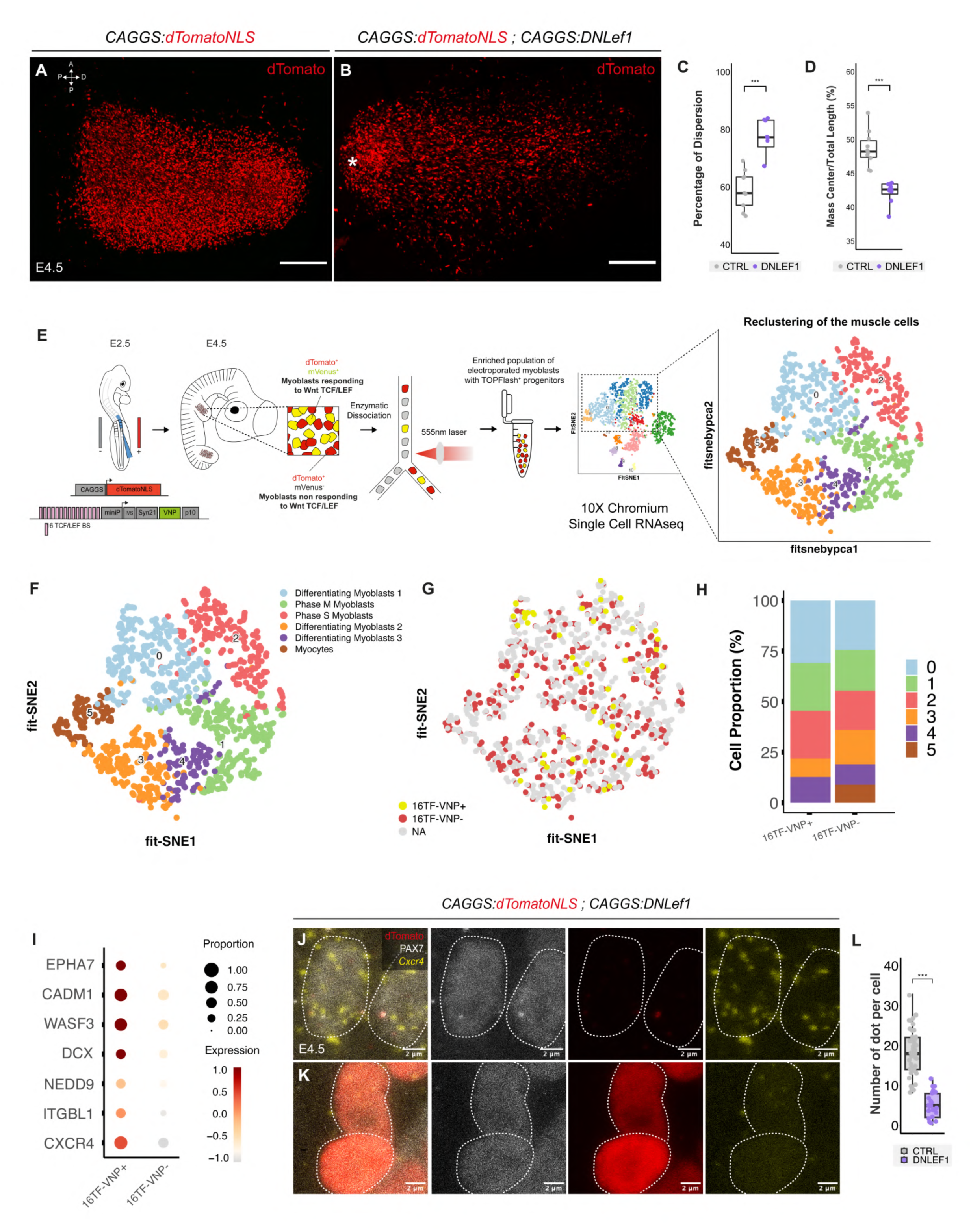
TCF-LEF/β-catenin controls the spatial distribution of primary myotube progenitors via CXCR4. (**A-B**) Dorsal view of a confocal stack of limbs electroporated with an ubiquitously expressed nuclear tomato alone (**A**) or together with DN Lef1 (**B**). (**C-D**) Quantifications of the repartition of muscle progenitors in the limb. (**E**) procedure followed to perform scRNAseq on electroporated limb muscle progenitors. (**F**) six distinct clusters were identified within the total population of myoblasts. (**G-H**) Tomato-positive cells (red) and venus-positive cells (yellow) distribute in all, but cluster 5. (**I**) List of migration-associated genes differentially expressed in the 16TF-VNP+ and 16TF-VNP-populations of myoblasts. (**J-K**) HCR FISH to detect CXCR4 expression in control limb myoblasts (**J**) or after DN Lef1 over-expression (**K**). (L) Quantification of (**J,K**).

Numerous Wnt ligands and Wnt antagonists are expressed during limb formation in amniote embryos ^73–75^, and some of these could potentially regulate the migration and/or the distribution of the primary myotube progenitors. An important question was whether Wnt signaling controls this process directly, acting as a trophic factor, or indirectly, through transcriptional regulation of another molecule. To test this, we injected Wnt-expressing cells into early limb buds at E3.5 and analyzed the distribution of (PAX7+) muscle progenitor cells one day after. These manipulations did not lead to detectable defects in the normal distribution of muscle progenitors in experimental embryos (Supplementary Figure 9A-C), suggesting that, while Wnt signaling regulates progenitor migration, Wnt ligands do not act as positional cues in this process.

A multitude of genes and signaling pathways are known to regulate muscle progenitor migration during embryogenesis ^76–79^, making a candidate gene approach to test targets of TCF-LEF/β-catenin signaling in primary myotube progenitors a tedious endeavor. Therefore, we adopted a transcriptomic approach, performed on limb progenitors co-electroporated with a ubiquitously-expressed nuclear dTomato and the 16TF-VNP reporter, followed by cell sorting based on red fluorescence. This enriched population of myoblasts was then subjected to single-cell RNA sequencing (scRNAseq) analysis (Figure 5E).

Using unsupervised graph-based clustering combined with FFT-accelerated interpolation-based t-SNE (Flt-SNE), we identified several distinct clusters of cells (Figure 5E). We employed negative selection to exclude non-muscle contaminating cells, identified by specific markers (e.g., CSF1R for macrophages; FLT for endothelial cells, see Supplementary Figure 10A), while retaining cells expressing PAX7, a known marker of muscle progenitors. Within PAX7+ cells, we identified six distinct clusters that, based on the expression of known factors (Supplementary Figure 10B,C), represent stages of proliferation and differentiation of muscle progenitors (Figure 5F). By mapping for the presence of transcripts from the two electroporated transgenes within these clusters, individual cells could be grouped into two distinct populations: 16TF-VNP- (corresponding to secondary myotube and satellite cell progenitors) and 16TF-VNP+ (encompassing progenitors of the primary myotube lineage; red and yellow dots, respectively, Figure 5G,H). The 16TF-VNP+ cells were absent in Cluster 5, which comprises progenitors in the late stages of myogenic differentiation (e.g., MyoD+, MyoG+, Mymk+; Supplementary Figure 10C), consistent with earlier data (Figure 2). However, cells from the two populations were found in all other clusters, suggesting that at that stage of development, their transcriptional signatures are largely similar.

Despite this high similarity, differential expression gene analysis between the 16TF-VNP+ and 16TF-VNP-populations revealed that i) known members and targets of Wnt canonical signaling were expressed at a higher level in the 16TF-VNP+ cell population, while secreted inhibitors of the pathway were more prevalent in the 16TF-VNP-cell population (Supplementary Figure 10D); ii) factors known to regulate cell migration in various contexts were overexpressed in the 16TF-VNP+ population (EPHA7 in the hematopoietic system ^80^; EPHA7 and DCX in the nervous system ^81,82^; CADM1, NEDD9, WASF3 and ITGBL1 in cancer ^83–86^; Figure 5I). These findings align with a role of TCF-LEF/β-catenin signaling within the 16TF-VNP+ population in the transcriptional regulation of a range of putative migration regulators. The chemokine receptor CXCR4 emerged as a particularly intriguing candidate. In mouse and chicken embryos, CXCR4, together with its cognate ligand SDF1, regulates muscle progenitor migration. CXCR4 is expressed in a subset of limb muscle progenitors, while SDF1, expressed in the limb mesenchyme, acts as a trophic factor to guide muscle progenitor migration ^87^.

These observations suggested that CXCR4 might be a target of TCF-LEF/β-catenin-dependent signaling in early limb progenitors of the primary myotube lineage. To test this hypothesis, we electroporated the DN Lef1 construct at E2.5 in the VLL of brachial somites. Two days later, the limb buds were analyzed by HCR FISH ^88^ to detect the expression of CXCR4. We observed that the expression of CXCR4 was strongly downregulated in electroporated PAX7+ myoblasts compared to the PAX7+ WT myoblasts (Figure 5J-L). Altogether these results strongly suggest that TCF-LEF/β-catenin-dependent signaling in the progenitors of the primary myotube lineage is essential to regulate their distribution in the growing limb bud, partially or entirely through the transcriptional regulation of the CXCR4 receptor.

## Discussion

The discoveries outlined in this study address two enduring puzzles that have persisted within the field of myogenesis for four decades.

i) Firstly, the study resolves the longstanding question of the developmental origin of primary and secondary myotubes. It unequivocally demonstrates that these myotubes originate from two distinct myogenic progenitor populations co-existing early in the limb, distinguished by their TCF-LEF/β-catenin-dependent Wnt signaling activity.
ii) Secondly, our finding clarifies a controversy surrounding the lineage association between embryonic and fetal myoblasts. The observation that myogenic precursors isolated from embryonic and fetal limbs exclusively generated primary or secondary myofibers, respectively, had led to the conclusion that myogenic progenitors consist of a homogeneous precursor population that evolves as development progresses ^27,29,30^. Contradictory observations showed that embryonic and fetal myoblasts are two distinct precursor populations, sequentially migrating into the limb mesenchyme ^33^, or present from the onset of limb formation ^10,36^. Our study resolves this controversy by demonstrating that the myogenic progenitors present during limb development comprise a mixed population of precursors for primary and secondary myotubes. A surprising outcome of our findings is the identification of a shared origin for limb secondary myotubes and satellite cells. Although it is long established that all limb muscles, including its satellite cell component, originate from the VLL ^89,90^, the developmental path that VLL cells follow to generate satellite cells, once they have migrated into the limb mesenchyme, had not been investigated.

These findings prompt a re-evaluation of muscle development in both trunk and limb. Trunk muscle formation has been thoroughly studied and is understood to occur in two distinct stages: i) an initial stage, where epithelial cells from the edge of the dermomyotome closest to the neural tube contribute to the formation of the primary myotome, composed of mononucleated myocytes ^5,7,91^. This primary myotome then transitions into a polynucleated structure through fusion with cells from other dermomyotome borders ^40^; ii) a second stage, where satellite cell progenitors emerge from the central dermomyotome and migrate into the primary myotome. These progenitors initially fuse with each other to form additional polynucleated fibers, with a subset set aside to become satellite cells ^6,60,61^.

The similarities between these processes in the trunk and our observations in the limb suggest a compelling hypothesis: despite the apparent differences in their origins—trunk muscles forming within the structured somite and limb muscles arising from migrating progenitors—both may follow a similar developmental pattern involving two distinct progenitor populations with different fates: i) the primary myotome and cells from the dermomyotome borders in the trunk may be analogous to the primary myotube lineage we identified in the limb; ii) cells from the central dermomyotome could correspond to the TCF-Trace-negative progenitors in the limb, along with their derivatives, the satellite cells.

Further analyses will be necessary to determine whether the biochemical and cellular characteristics of these myogenic progenitor populations support a unified model of trunk and limb muscle formation.

As was observed in the trunk ^40^, it is fascinating that limb muscle progenitors carefully choose their fusion partners in an environment where all progenitors and myofibers are tightly intertwined, such that TCF-Trace^+^ progenitors only fuse to myofibers derived from their own lineage, while TCF-Trace^-^ progenitors, at least initially, fuse to myofibers from their own lineage. The cellular and molecular underpinning of such intricate fusion pattern remains to be uncovered.

Lastly, the use of the TCF-LEF/β-catenin-dependent signaling reporter was pivotal in revealing the embryonic origins of two distinct populations of myogenic progenitors and their derivatives. This prompted a deeper investigation into the role of this pathway in limb myogenesis, leading to the discovery of its involvement in the spatial distribution of primary myotube progenitors. The identification of CXCR4, a known regulator of limb progenitor migration, as a transcriptional target of TCF-LEF/β-catenin-dependent signaling, underscores Wnt canonical signaling’s role in migration, likely acting indirectly through CXCR4. However, the identification of additional transcriptional targets associated with cell migration across various contexts suggests that CXCR4 may only partially mediate this role in the limb.

This finding adds yet another layer to the diverse functions of TCF-LEF/β-catenin-dependent signaling during myogenesis, complementing the extensive body of research on Wnt signaling in this process (see ^64,65,92^ for reviews). While many studies have suggested an essential role for Wnt signaling in the cell decision-making to initiate myogenesis in somites, the intricate morphogenetic processes that accompany this cell fate decision—such as the epithelial-mesenchymal transition of dermomyotomal cells, their migration into the primary myotome, and their orientation and growth along the antero-posterior axis—have made it challenging to pinpoint the specific roles of canonical Wnt signaling. This complexity might explain the contradictory findings in the field ^3,66,67,93–95^.

Divergent data also exist regarding Wnt signaling’s role in limb myogenesis and adult muscle regeneration, with studies suggesting that Wnts can promote, inhibit, or be dispensable for myogenesis ^69,96,97^. Given the extensive literature on Wnts’ potential roles in myogenesis and the numerous Wnts expressed during limb development ^74,75,98^, it was surprising to observe that the 16TF-VNP reporter activity was restricted to a relatively short time window of limb myogenesis. It was completely absent in one lineage and silent during the majority of late myogenesis, including in satellite cells. These observations rule out the possibility that Wnts, via canonical, TCF-LEF/β-catenin-dependent signaling, play a role in late myogenesis, tissue patterning, or the emergence of the satellite cell population.

Our findings align with previous research using genetic approaches in mice, which showed that the ablation of β-catenin in limb muscle progenitors does not affect MYOD expression and myofiber formation early (at E12.5) during embryogenesis ^55^. That study also suggested that there exists a PAX7+ lineage that contributes to late myotube formation. However, this study did not analyze the significance of those findings the context of the primary and secondary myotube lineages.

The discovery of a molecular signature distinguishing primary from secondary myotube lineages is poised to revitalize a field that has been largely dormant for years due to the absence of specific molecular markers for these lineages. This breakthrough opens new avenues for investigation, enabling comprehensive molecular characterizations of both lineages. This will allow us to address critical questions about the developmental pathways their progenitors follow to generate functional muscles in the embryo and their roles during development, as well as during regeneration and repair in the adult. The 16TF-VNP reporter and the TCF-Trace system, combined with advanced "omics" technologies, will be instrumental in deciphering the molecular pathways involved in some of the most enigmatic events of late myogenesis.

## Methods

### *In ovo* chicken embryos electroporation

Fertilized chicken eggs were obtained from a local breeder. To target wing muscle progenitors, the lateral borders (referred to as the VLL) of brachial somites in E2.5 chicken embryos were electroporated as described previously ^40^. Briefly, chicken embryos were incubated at 37.5°C until stage HH16 (52 hours or E2.5). The plasmid solution was injected into the brachial somites (somites 17-21) using a glass capillary. Three pulses of 50V, 10ms in duration, and spaced by 10ms, were applied directly to the embryo using tungsten and platinum electrodes. Eggs were then sealed and placed back in the incubator until the desired stage was reached. Cells emanating from the VLL and migrating into the wing bud mesenchyme constitute a mixed population of very early migrating endothelial progenitors, followed by myogenic progenitors^39,89,99–101^. This migration occurs over a short window of time, beginning soon after somite formation and lasting 15-20 hours ^89,99^. We developed an electroporation protocol that primarily targets myogenic progenitors (Supplementary Figure 1A).

### Expression constructs

All constructs used in this study contained two Tol2 sequences (T2: transposable elements from medaka fish ^38^) surrounding the entire construct to allow their stable integration in the chicken genome in the presence of transposase, provided by a co-electroporated transposase plasmid (CAGGS-Transposase). This plasmid does not contain Tol2 sequences and is thus gradually diluted along with cell division.

#### Electroporation marker

Depending on the experimental or immunostaining strategies, several plasmids were used to visualize electroporated cells. These plasmids contained the coding sequences for various fluorescent proteins with different spectral characteristics and/or directed to different cellular compartments. The T2-CAGGS:BFP and T2-CAGGS:mVenus plasmids were used to label the cytoplasm of all electroporated cells with blue and green fluorescent proteins, respectively. The T2-CAGGS:H2B-BFP, T2-CAGGS:H2B-Achilles and T2-CAGGS:dTomatoNLS were used to labelled the nuclei of all electroporated cells with blue, green and red fluorescent proteins, respectively. H2B- and NLS-fused fluorescent proteins were used to label single cells.

#### Wnt canonical signaling

Wnt binding to its receptor can trigger three downstream pathways. The best characterized (and referred to as ‘‘canonical’’) Wnt cellular response results in the inhibition of the β-catenin destruction complex, ultimately leading to the translocation of β-catenin into the nucleus where it partners with members of the TCF/LEF family of transcription factors to activate various Wnt target genes. To monitor this canonical transcriptional response, efficient transcription-based reporter systems that combine the DNA binding sites of TCF/LEF upstream of a minimal promoter and a reporter gene have been designed, the first of which named TOPFlash ^102^. Since the first description of the TOPFlash reporter, many derivatives have been generated that combined multiple TCF/LEF binding sites driving a variety of reporter genes. The high stability of the most commonly used reporters (β-galactosidase, GFP, RFP …) leads to their considerable accumulation in cells activating the pathway, thus greatly facilitating their detection. However, two major drawbacks are: i) an important lag-time between the activation of the pathway and the accumulation of sufficient amounts of the reporter to be detected and ii) conversely, the detection of signals in tissues where the activity of the pathway may have already ceased. Most of these constructs are therefore unsuitable to detect rapid spatiotemporal changes in TCF-LEF/β-catenin dependent signaling. We have used here a recently developed reporter of Wnt/β-catenin dependent signaling activity, named 16TF-VNP ^37^, where the combination of 16 TCF/LEF-binding sites ^102^, translational enhancers ^48,103^, and a destabilized (half-life, 1.8 hours ^104^), fast maturing Venus fluorescent protein ^105^ generated the most sensitive to date, yet dynamic, molecular tool to detect TCF/β-catenin-responding cells and tissues in developing embryos, with no detectable background ^37,106^.

#### Short-term lineage tracing

For short-term pseudo-lineage of TCF/LEF responding cells, we inserted a P2A sequence downstream of and in-frame with the 16XTF-VNP reporter, followed by a stable form of nuclear mCherry (half-life, about 18 hours). This results in the production of two fluorescent proteins with different half-life. This strategy was described previously in *Drosophilia* ^48^ to generate a dual-color Trans Timer construct that provides spatio-temporal information on signaling pathway activities.

#### Long-term lineage tracing

Even stable fluorescent proteins, such as GFP or RFP, are not suitable for long-term studies. For this, we have engineered a tri-partite lineage tracing system based on the Tet-On and the Cre-Lox technologies. A plasmid containing the destabilized rtTA transactivator gene under the control of the 16TF promoter was mixed with a plasmid encompassing a bi-directional TRE (Tetracycline Response Element) promoter (pBI) driving the expression of a CRE recombinase (T2-pBI:CRE) and a plasmid coding for a CRE-inducible nuclear dTomato (T2-CAGGS:LoxP-STOP-LoxP-dTomatoNLS) or a cytoplasmic eGFP (T2-CAGGS:LoxP-STOP-LoxP-eGFP).

As the electroporation technique results in the mosaic expression of transgenes in targeted cells, a ubiquitously-expressed electroporation marker was added to the plasmid mixes to identify and analyze all electroporated cells.

### Wholemount immunohistochemistry

E3 to E7.5 electroporated limb buds were imaged in whole mount as described ^40^. Briefly, after dissection, samples were fixed for 1 hour in 4% formaldehyde/PBS at room temperature (RT), briefly rinsed with PBS, pre-incubated in washing buffer (0.2% BSA, 0.1% Triton X-100, 0.2% SDS in PBS) for 1 hour at RT, and incubated overnight at 4°C with the primary antibodies. They were washed at least 5 times during the following day and incubated with secondary antibodies overnight at 4°C. As fixation significantly decreases the brightness of fluorescent proteins (particularly GFP), we often used primary antibodies against fluorescent proteins to facilitate their detection. The following antibodies were used: rabbit polyclonal antibody (IgG) against dTomato (ab62341, Abcam, 1/1000); chicken polyclonal antibody (IgY) against eGFP, mVenus and Achilles (A10262, Invitrogen, 1/1000); mouse monoclonal antibody (IgG2a) against eGFP, mVenus and Achilles (A11120, Invitrogen, 1/1000); rabbit polyclonal antibody (IgG) against TagBFP (AB233, Evrogen, 1/500); mouse monoclonal antibody (IgG1) against PAX7 (AB528428, DSHB, 1/10); mouse monoclonal antibody (IgG2b) against Myosin Heavy Chain (MF20, AB2147781, DSHB, 1/10) and mouse monoclonal antibody (IgA) against the slow myosin MYH7B (S58, AB528377, DSHB, 1/10). These primary antibodies were used in combination with species-matched secondary antibodies conjugated with Alexa fluorochromes (488nm, 555nm or 647nm) from ThermoFisher or SouthernBiotech at a concentration of 1/500. After staining, samples were incubated in 50% glycerol/PBS for 1h at RT and in 80% glycerol/PBS for several hours before analysis.

### Cryosectionning

E9.5 to E18.5 limb buds were fixed for whole-mount imaging as described, washed in PBS for 1 hour, and then incubated in 7.5% sucrose/PBS and 15% sucrose/PBS at room temperature (RT) for 30 minutes to several hours, depending on their size. Next, they were immersed in a pre-melted solution of 15% sucrose and 7.5% bovine skin gelatin at 42°C with agitation overnight before being placed into cryosectioning molds. The samples were then immersed in dry ice-cold 100% ethanol and stored at - 80°C until sectioning. They were cryosectioned using a Leica cryostat at a thickness of 18 µm. Sections were stored at −80°C until analysis. Immuno-histochemistry was performed as described for whole-mount imaging, using the same set of antibodies, with the exception that secondary antibodies were incubated for only 2 hours at RT. Nuclei were detected using DAPI. The slides were mounted in Fluoromount ^TM^. Transversal and longitudinal sections were performed for the analyses of the 16xTF-VNP reporter at E9.5, while only transversal sections were done at E12.5, E14.5, E16.5 and E18.5. Analysis of the TCF-Trace experiments after E9.5 were performed with transversal sections targeting the extensor muscle of the zeugopod.

### Doxyxycline induction

Stock solution of doxycycline at a concentration of 20mg/ml in ddH2O was prepared in advance and stored at −20°C. A solution at 3,5μg/ml was prepared by diluting the stock solution into sterile Ringer’s solution on the day of the injection; 300μl of the solution was added per embryo.

### EdU incorporation

50μl of 10mM of EdU solution was added directly onto the embryo that was placed back in the incubator for 1h. Embryos were then dissected, fixed and immunostained as described above. Once immunostained, samples were pre-incubated in 250μl of PBS with 1μl of Alexa fluorophore for 1h at RT. Separately 150μl of PBS was mixed with 100μl of ascorbic acid at 0,5M and 2μl of a 1M CuSO4 solution and added to the pre-incubated samples. Embryos were incubated overnight at 4°C with agitation, washed at least five times the following day and cleared into glycerol as described.

### Imaging, quantifications and statistical analyses

For whole-mount and cryosectioned samples, imaging was performed using a Leica SP5 confocal microscope with a resolution of 1024×1024 pixels, utilizing either a 20x or 40x objective with a Z-step size of 2 µm. Images were analyzed using Fiji with the Cell Counter plugin. Plot and statistical analyses were done using R, with ggplot2 and ggsignif packages. Experiment with two different conditions were compared using Wilcoxon-Mann-Whitney test and experiment with three different conditions with Kruskal-Wallis test associated with Dunn-Bonferroni post-hoc test. NS represent a p-value>0.05, *** a p-value<0.001 and **** a p-value<0.0001

### Purification of electroporated single cell

E4.5 electroporated chicken embryos with a ubiquitous dTomato and the 16TF-VNP reporter were screen under a fluorescent binocular and electroporated limb buds were quickly dissected and incubated with 500 μl of pre-warmed Dispase (1,5mg/ml in DMEM / 10mM Hepes), pipette up and down 10 times and incubated 15min at 37°C. The sample was homogenized every 5min then 500 μl of pre-warmed Trypsin (0,05% in DMEM) was added to the tube, homogenized and incubated 3min at 37°C. Samples were then transferred into a 15ml falcon tube, and the reaction was stop with 10ml of Hanks buffer (for 100ml: 10ml of HBSS 10X, 250mg of BSA, 1ml of Hepes 1M, in sterile ddH_2_O), homogenized and centrifugated 10min at 500g. The pellet was re-suspended in 4ml of Hanks buffer and filtered with a pre-humidified 40 μm sterile filter and re-centrifugated 10min at 500g. The final pellet was re-suspended into 250 μl of Hanks buffer and added to 250 μl of Hanks buffer in a pre-humidified FACS tube. For sorting, we added DAPI (1/1000) in the final Hanks buffer solution and prepare a sample containing non-electroporated tissues, and non-electroporated tissues stained with DAPI to calibrate the sorting. Cells were then sorting according to the dTomato fluorescence and collected into Hank’s buffer. For the single-cell RNA-seq experiment, a total of 6 electroporated limb buds were pooled together in the same tube.

### Single cell RNA-seq analyses

To profile cell-type composition after electroporation, we performed single cell RNA-sequencing (snRNAseq) on chicken embryo limb bud electroporated with the CAGGS:dTomatoNLS and the 16TF:VNP plasmid (see section above). Data demultiplexed in fastq format was checked with fastqc and fastq_screen to control quality sequencing. Sequencing reads were processed with STAR (v.2.7.9) to align and quantify data. Genome and transcriptome used is GRCg6a.105 from Ensembl database & custom references gtf and fasta files corresponding to the plasmids used. We merged both datasets in one. The downstream treatment for analysing separately and integratively the data after filtering was *SCTransform* with *glmGamPoi* method and a regression of percentage of mitochrondrial, PCA on SCT assay. We determined Doublet barcode with *scDblFinder* R packages on cluster-based approach and *dbr=0.1*. We selected any features where mean (count _sample_)_feature_ > 0.1. (s)KNN graph build on pca reduction matrix with a dimension depending of sample (we selected the last elbow point where cumulative sums of percentage Standard Deviation > 90% and the n+1 dimension not exceed < 1% supplementary), we used Louvain Clustering with *igraph* method. We used also FFT-accelerated Interpolation-based t-SNE (FItSNE) with these parameters : a modification of initialization (first and second PCA dimensions are divided by standard deviation of first PCA dimension multiplied by 0.0001), *perplexity_list* used is a list starting 30 to nCells_integrated dataset_/100) and *learning rate = nCells_integrated dataset_ / 12,* all PCA dimensions selected from the last step and 1000 iterations to run. Biomarkers are identified after *PrepSCTFindMarkers* on SCT assay and performed with *FindAllMarkers* with MAST test. We filtered filtered biomarkers to conserve only features |pct.1_feature/cluster_ – pct.2_feature/cluster_ | > 0.2 and p-adj < 0.05. We selected a top5 on each cluster found and sign of log2FC. In total, 2108 cells were sequenced and used for further analysis. We identified the presence of absence of the dTomato and the mVenus transcripts and identified 432 cells where dTomato only was expressed ≥ 15 (∼19%) & 55 cells where mVenus was expressed ≥ 15 both (∼2.6%). Each cell covered a value range of nCount RNA between 805 to 20000, nFeature RNA between 601 to 4462 and %mitochondrial ≤ 0.7 %. Muscle progenitors dataset was isolated after identification of clusters based of genes expression and the same protocol of downstream refinement is used (SCT + PCA + dynamic selection of dimension PCA’s).

### Analysis of DN Lef1 phenotypes

Images of E4.5 control and DN Lef1 limb bud were acquires in wholemount on the confocal as described above. 3D acquisitions were transformed into 2D images with a maximal projection in Fiji. For the percentage of dispersion, an oblong shape was drawn around the muscle mass to measure its total area and then a threshold was set to measure the area of all the nuclei using the “analyze particles” function. The percentage of dispersion was calculated as follow:

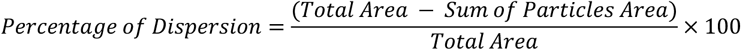

For the center of mass, a line was traced in the middle of the muscle mass, along the proximo-distal axis of the limb and the center of mass was measured using the “Measure” function. The center of mass is defined as the brightness-weighted average of the x and y coordinates of all pixels in the image or selection. Therefore, if the repartition of the fluorescence is homogeneous along the proximo-distal axis the center of mass should be around the middle of the region of interest. The center of mass was normalized by the total length of the muscle mass, therefore a value around 50 represent a homogeneous repartition.

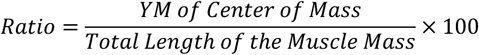

### In ovo cell grafting

Control NIH 3T3 or Wnt3a secreting CRL-2647 cells were grown in DMEM / 1% Penicillin-Streptomycin / 10% FBS until confluence. On the day of injection, cells from two 150mm plates for each condition were collected after addition of 3ml of 0,25% trysin in DMEM. The reaction was blocked with DMEM / 1% Penicillin-Streptomycin / 10% FBS and cells were harvested in a 15ml Falcon tube, centrifuged and washed twice with 10ml of PBS. In a separated tube, 1μl of CFSE CellTrace™ (#C34570, ThermoFisher) was mixed with 700μl of PBS. Each pellet of cells was homogenized with 300μl of the CFSE mix and incubated 20min at 37°C with regular homogenization to avoid cell sedimentation. The reaction was blocked with DMEM / 1% Penicillin-Streptomycin / 10% FBS, cells were centrifuged and supernatant removed to obtain a thick preparation of cells. Cells were then loaded into a glass capillary (30-0047, Harvard Apparatus) with a Hamilton Microliter™ syringe #805 and injected with a PicoSpritzer injector into E3.5 chicken embryo limb buds.

## Supplementary Figures

**Supplementary Figure 1: Early expression of the 16TF-VNP reporter in muscle progenitors. (A,B)** Dorsal view of a confocal stack of E3 brachial somites electroporated with an ubiquitous dTomato (in red), the 16TF-VNP reporter (in green) and stained for PAX7 (in grey). **(B)** is an enlargement of **(A)**, arrowheads indicate electroporated cells that are 16TF-VNP^-^, but positive for PAX7. **(C)** Quantification of the percentage of PAX7^+^ cells that are 16TF-VNP^-^ and 16TF-VNP^+^. **(D)** Longitudinal section of E4.5 limb electroporated with an ubiquitous dTomato and the 16TF-VNP reporter. Dotted line indicate the junction between the dermis and the epidermis. **(E)** Longitudinal optical section of limb dorsal muscle mass electroporated with an ubiquitous TagBFP (in blue) and the 16TF-VNP reporter (in green). **(F)** Ventral view of a confocal stack of the ventral muscle mass of a E4.5 limb bud electroporated with a ubiquitous TagBFP and the 16TF-VNP reporter. Scale bars: 50μm (B), or 100μm (A, D-F)

**Supplementary Figure 2: The 16TF-VNP reporter is not active in late embryonic muscle masses (A-E)** Longitudinal sections of E9.5 limb buds electroporated with an ubiquitous nuclear dTomato (in red), the 16TF-VNP reporter (in green) and stained for DAPI (in blue) and PAX7 in grey) at various anatomical locations along the limb proximo-distal axis. Scale bars: 100μm (A-E)

**Supplementary Figure 3: The 16TF-VNP reporter is not active in muscle masses in fetal stages. (A-H)** Transversal sections of E12.5 **(A,B),** E14.5 **(C,D),** E16.5 **(E,F))** and E18.5 **(G,H)** limb buds electroporated with an ubiquitous nuclear dTomato (in red), the 16TF-VNP reporter (in green and stained for DAPI (in blue) and PAX7 (in grey). **(B), (D), (F), (H)** are enlargements of **(A), (C), (E), (G)** respectively. Scale bars: 50μm.

**Supplementary Figure 4: PAX3 and PAX7 are co-expressed in muscle progenitors throughout development. (A)** Dorsal view of a confocal stack of a E4.5 limb bud electroporated with a ubiquitous TagBFP (in blue), the 16TF-VNP reporter (in green) and stained for PAX3 (in magenta) and PAX7 (in red). White arrowheads indicate 16TF-VNP^-^ cells, yellow arrowheads indicate 16TF-VNP^+^ cells. **(B)** Quantification of the percentage of PAX3+ cells within the 16TF-VNP^-^ and 16TF-VNP^+^ populations. Quantification of the percentage of PAX7+ cells within the 16TF-VNP^-^ and 16TF-VNP^+^ populations. **(D-G)** Transversal sections of limb muscles from E10.5 **(D),** E12.5 **(E),** E14.5 **(F)** and E16.5 **(G)** embryos stained for DAPI (in blue), PAX3 (in magenta) and PAX7 (in green). Each dot represents a limb bud. Scale bars: 50μm.

**Supplementary Figure 5: 16TF-VNP^+^ and 16TF-VNP^-^ progenitors proliferate at the same rate. (A-B)** Dorsal view of a confocal stack of E4.5 limb buds electroporated with a ubiquitous nuclear dTomato (in red), the 16TF-VNP reporter (in green) and labeled with EdU (in grey) to detect proliferating cells. White arrowheads indicate EdU^+^ cells; yellow arrowhead indicates EdU^-^ cells. **(C)** quantification of EdU^+^, 16TF-VNP^+^ or 16TF-VNP^-^ progenitors. Each dot represents an analyzed limb bud. Scale bar: 50μm

**Supplementary Figure 6: Pseudo-lineage of 16TF-VNP+ cells. (A,B)** Dorsal view of a confocal stack of E4.5 limb bud electroporated with a ubiquitous TagBFP (in blue) and the 16TF-VNP-P2A-mCherryNLS construct. The unstable VNP is detected in green; the stable mCherry is detected in red. **(B)** is an enlargement of **(A). (C)** Quantification (in percentage) of progenitors positive or negative for Venus and mCherry. Each dot represents an analyzed limb bud. Scale bars: 50μm (B) or 100μm (A)

**Supplementary Figure 7. Expression of slow myosin throughout embryonic and fetal limb bud development (A-H)** Transversal sections of whole limb bud from E7.5 to E18.5 stained for DAPI (in blue), MYH1 (Myosin Heavy Chain, in red) and MYH7 (Slow Myosin, in grey). **(I-P)** enlargements of limb bud muscles from E12.5 to E18.5 showing primary myotubes (MYH7^+^, white arrowheads) surrounded by smaller presumably secondary myotubes (MYH7^-^, yellow arrowheads). Scale bars: 10μm (J,L,N,P), 20μm (I,K,M,O) or 100μm (A-H).

**Supplementary Figure 8. Inhibition of TCF-LEF/β-catenin does not affect myoblast proliferation or myogenic differentiation, but regulates their spatial distribution.** (**A-B**) Dorsal view of a confocal stack of E4.5 limb buds electroporated with an ubiquitous nuclear dTomato (in red), alone (**A,C**) or together with an ubiquitously expressed DN Lef1 (**B,D**), stained for Phospho-histone H3 (in green, **A-B**) or EdU (in green, **C-D**). (**E-F)** are quantifications of (**A-D**). (**G-H**) Dorsal view of a confocal stack of E6.5 limb buds electroporated with by ubiquitous nuclear dTomato (in red) alone (**G**), or together with an ubiquitous DN Lef1 (**H**) and stained for MYF5. (**I**) are quantifications of (**G-H**). (K-L; N-O) are confocal stacks of images of limb buds electroporated with an ubiquitous nuclear dTomato (in red), together with a Tet-on inducible DN Lef1. IHC for MYOG (magenta, **K-L**). (**M**) quantification of (**K-L**). (**J**) depicts the experimental protocol used in (**K-O**) where DN Lef1 expression is activated by addition of doxycyclin.

**Supplementary Figure 9. Wnt does not play a trophic role on the migration of limb myoblasts. (A)** Control fibroblasts (NIH 3T3) or Wnt3a secreting cells (CRL-2647) were cultivated *in vitro*, labelled with green fluorescent CFSE and injected into the forelimb of a E3.5 chicken embryo and analyzed one day later. (**B,C**) Dorsal view of a E4.5 embryo after grafting of NIH 3T3 cells (**B**) or WNT3a/CRL-2647 (**C**) and immunostained for PAX7.

**Supplementary Figure 10. Single-cell transcriptomic analysis of sorted myoblasts.** (**A**) Expression of various markers for different cell lineages in raw dataset. (**B**) Heatmap showing the expression of some differentiated genes in the 6 myogenic clusters (**C**) Representation of some key genes in the myogenic clusters (**D**) Differential gene expression of Wnt-TCF/LEF related genes in the dTomato+ only and the dTomato+/mVenus+ population.

## Acknowledgements

We thank Drs Frank Stockdale, Frédéric Relaix, and Jean-Louis Bessereau for critical reading of the manuscript and insightful discussions on this study. We thank the Centre d’Imagerie Quantitative Lyon-Est (CIQLE) for imaging support. This research has been supported by a grant from the Agence Nationale de la Recherche (ANR). GT has received a PhD scholarship from the Ecole Normale Supérieure de Lyon (CDSN).

