## Supplementary figures and images for "Early lineage segregation of primary myotubes from secondary myotubes and adult muscle stem cells"

### Supplementary Figures 1-10

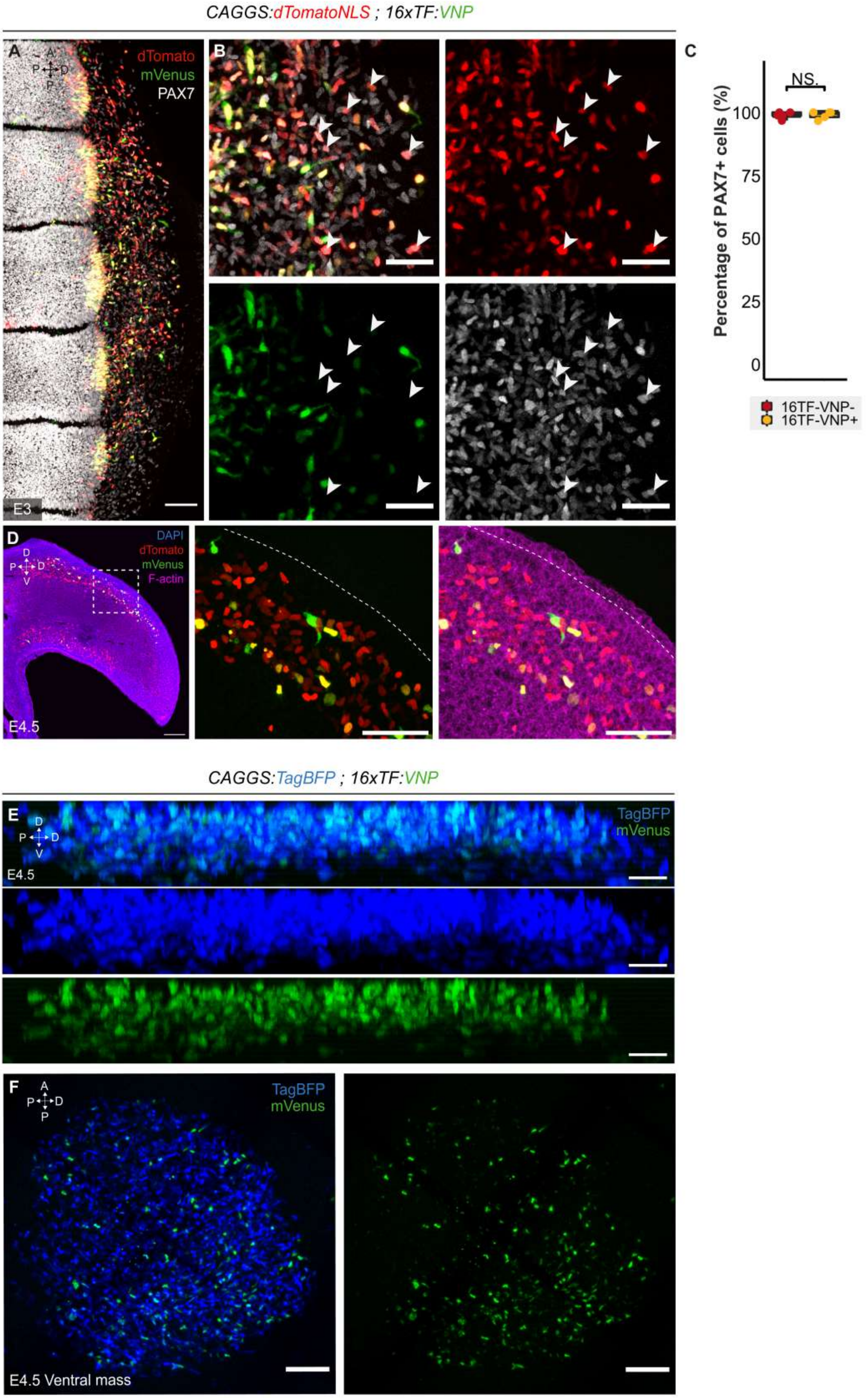

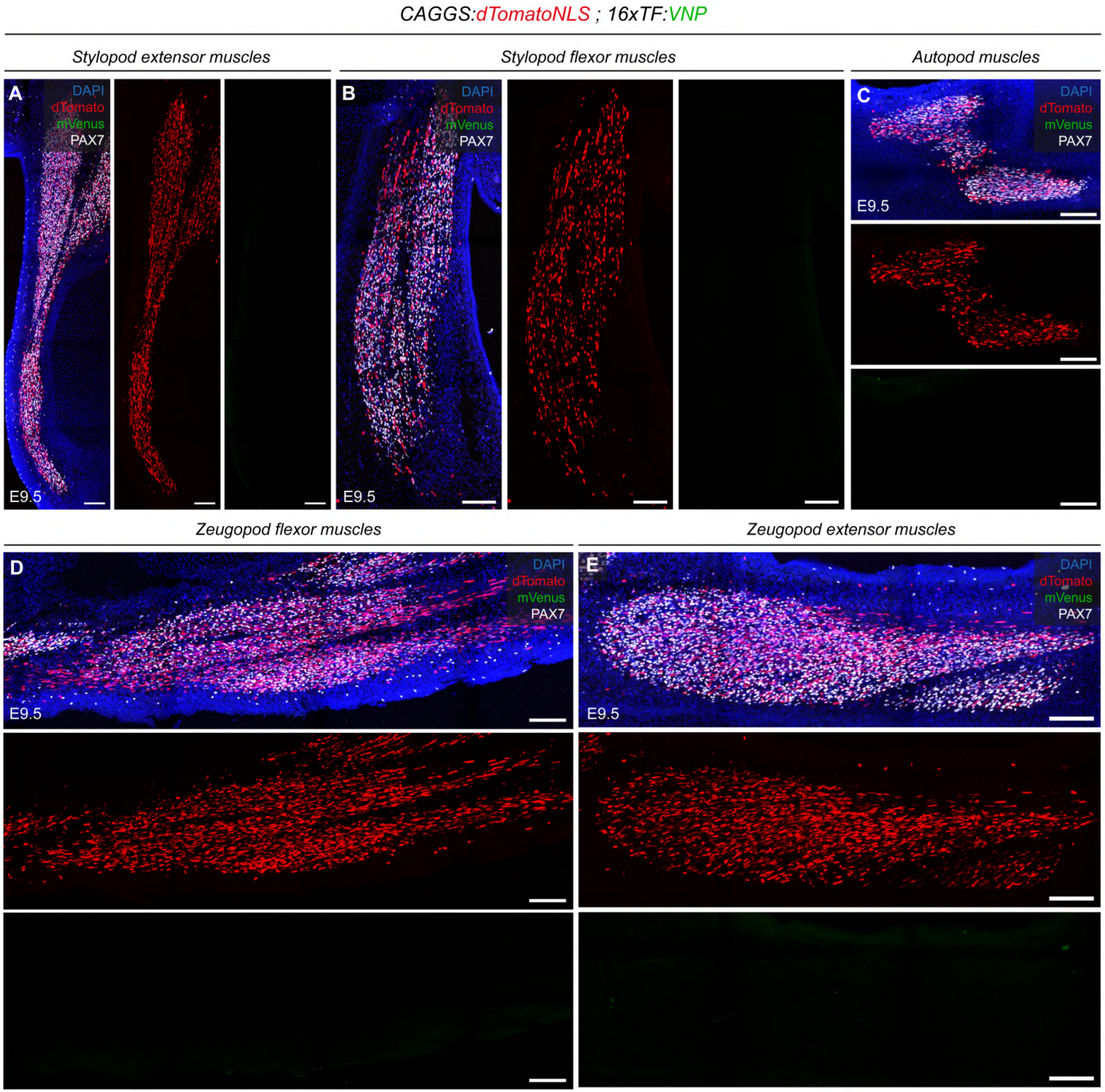

*CAGGS:dTomatoNLS ; 16xTF:VNP*

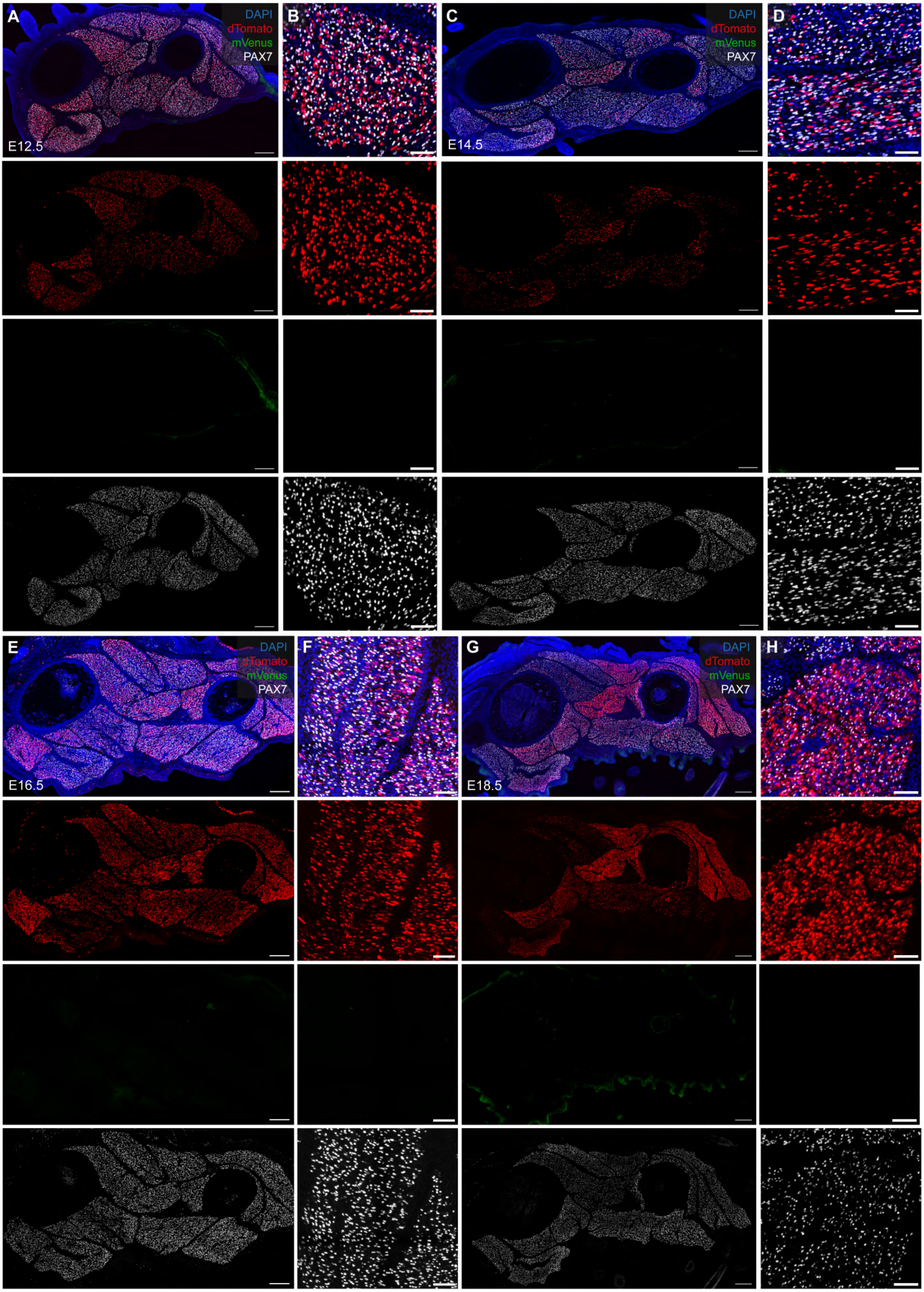

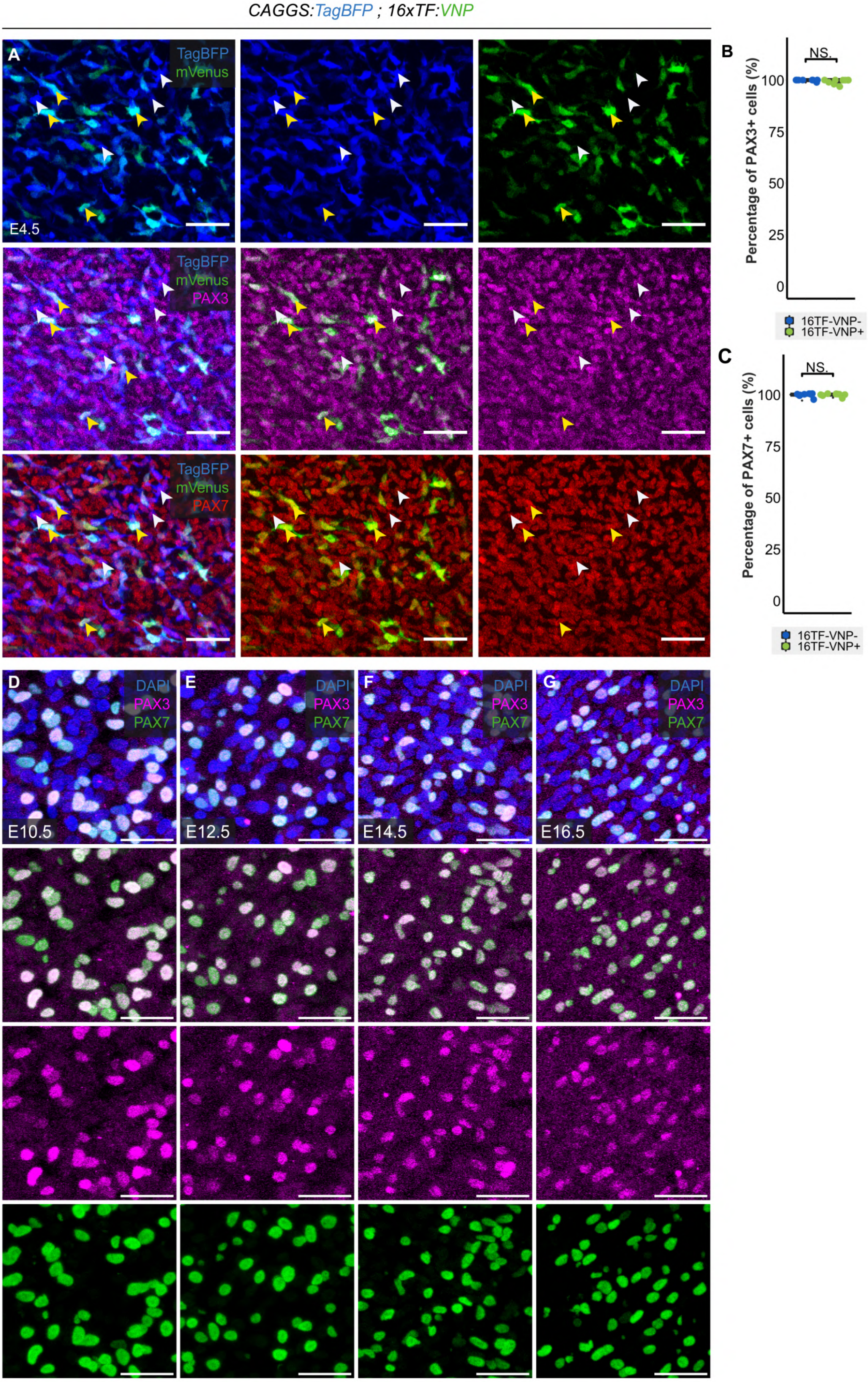

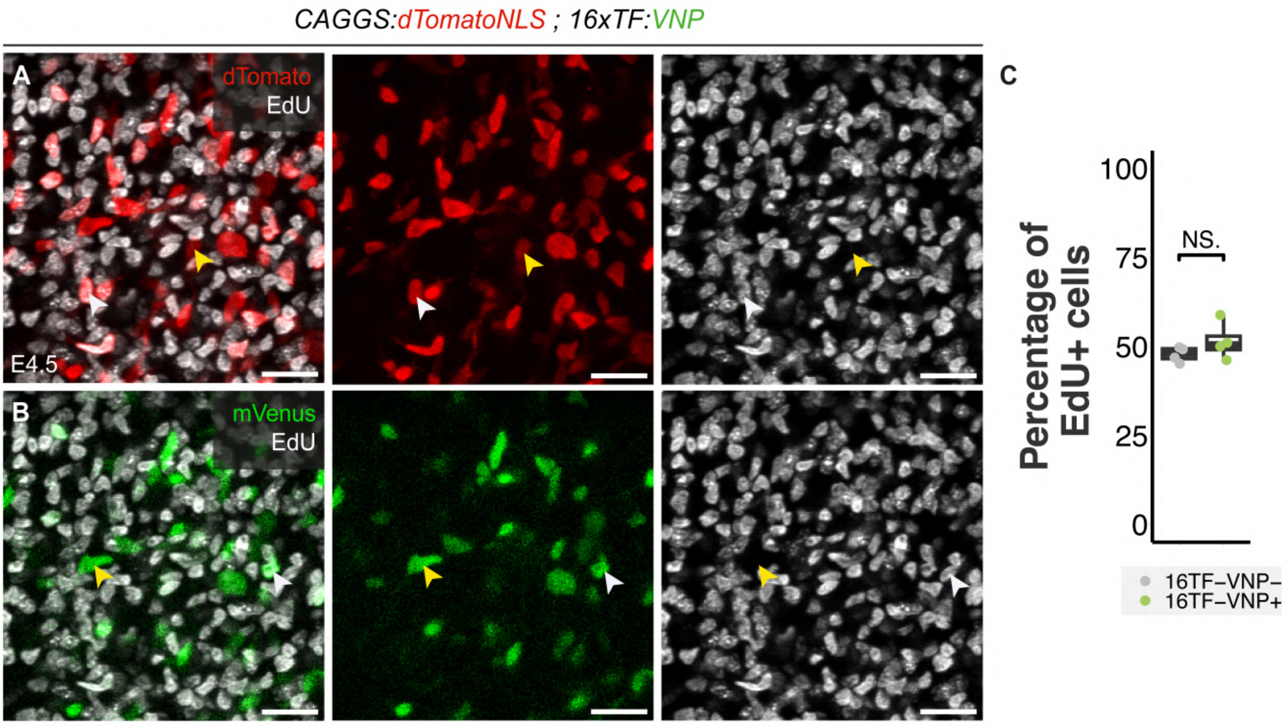

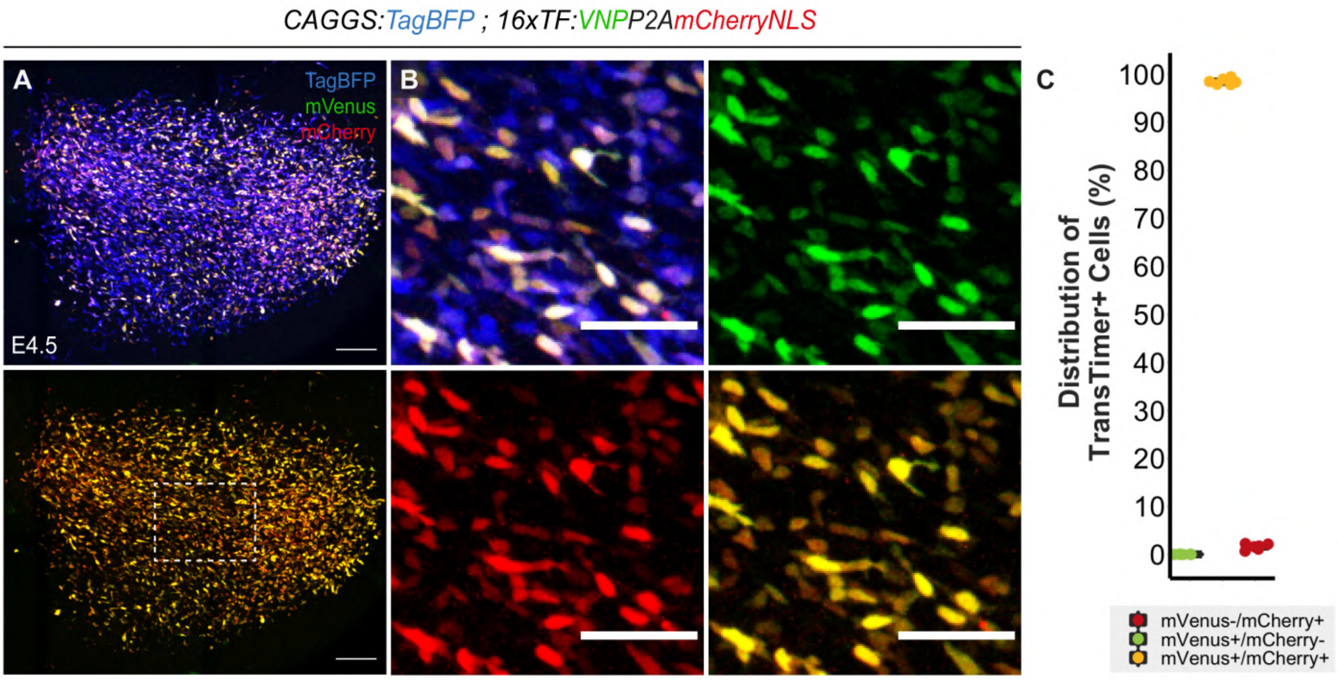

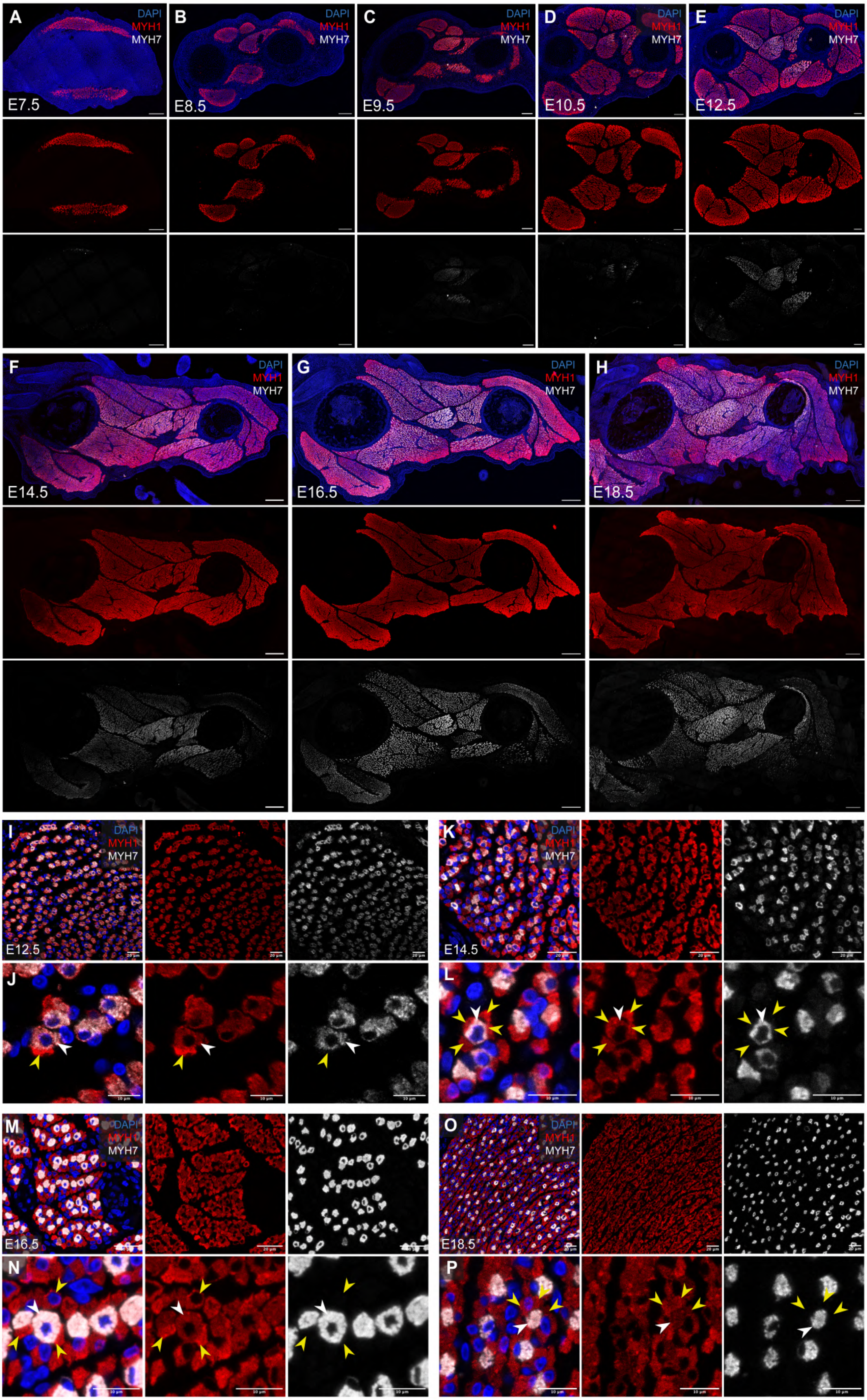

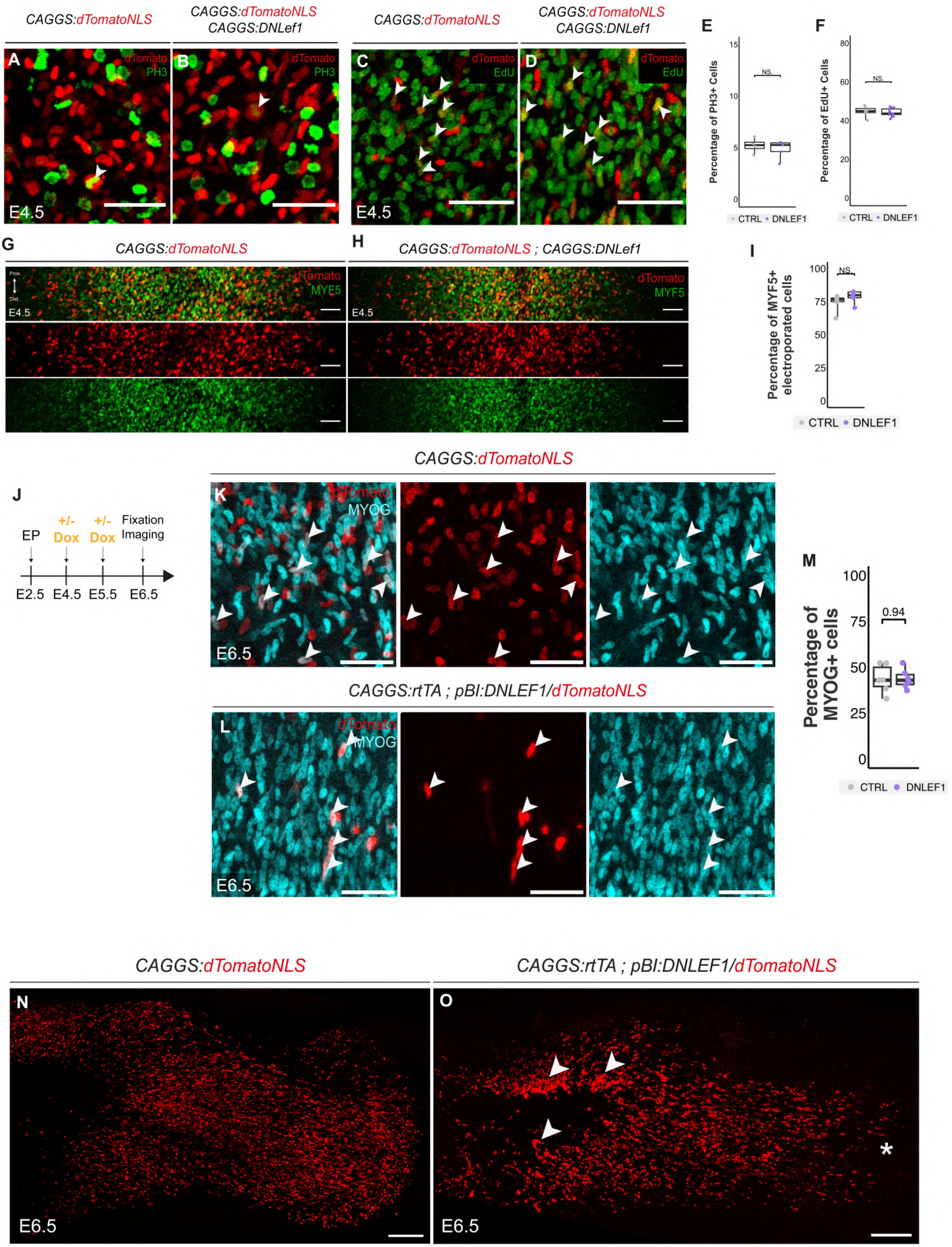

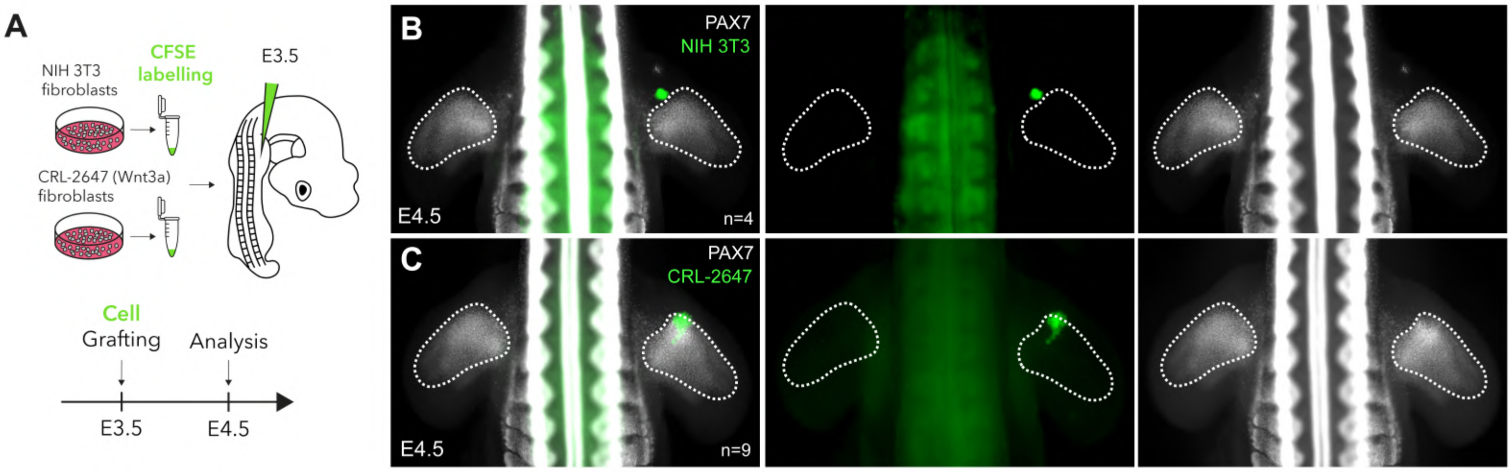

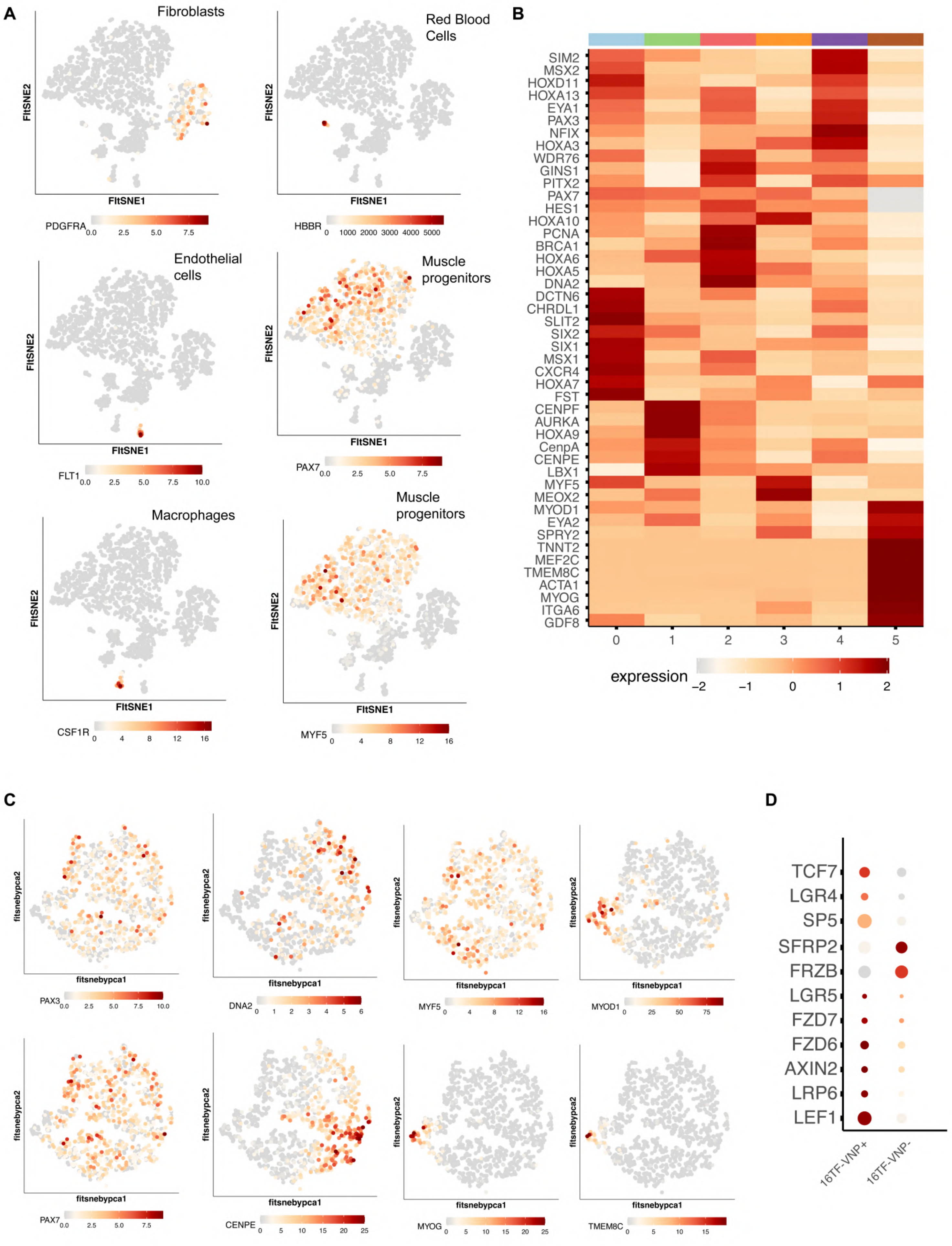
